# Nascent transcripts of LL2R tandem repeat nucleate locus-specific RNP-condensates recruiting splicing factors

**DOI:** 10.64898/2026.03.19.712899

**Authors:** Alla Krasikova, Anna Zlotina, Tatiana Kulikova, Veit Schubert, Anton Fedorov

## Abstract

Spatial organization of the cell nucleus is increasingly understood to be governed by non-coding architectural RNAs that seed the formation of biomolecular condensates. However, the genomic origins and mechanistic principles by which specific architectural RNAs orchestrate ribonucleoprotein (RNP) compartments remain poorly defined. Here, we characterised a novel non-coding architectural RNA that nucleates the formation of a locus-associated membraneless nuclear compartment. Using a combination of T2T genomic, transcriptomic, and high-resolution mapping on giant lampbrush chromosomes, we identified an array of non-coding LL2R tandem repeats on the long arm of chicken chromosome 2. The LL2R tandem repeat array is slowly transcribed by a convoy of RNA polymerases II as a single transcription unit from a retrotransposon promoter. With the use of scanning electron and super-resolution microscopy for detailed analysis, we demonstrate that nascent LL2R repeat transcripts are retained at the site of their transcription and recruit multiple sets of core splicing factors, including TMG-capped snRNAs, core snRNP proteins, and serine/arginine-rich proteins SRRM2 and SC35 that contain intrinsically disordered regions, thus seeding nuclear domain formation. At the same time, the RNP-matrix of the LL2R repeat transcription unit lacks hnRNP L, hnRNP K, and SFPQ, widely distributed along normal transcription loops that bear nascent gene transcripts. In contrast to pre-mRNAs, strand-specific LL2R tandem repeat-containing RNA contains more than threefold excess of predicted donor splice sites compared to acceptor splice sites and is depleted in polyadenylation signals. We suggest that stalling of elongating RNA polymerases at LL2R transcription units and delay in 3’ end cleavage may be caused by a deficiency of co-transcriptional splicing of nascent LL2R repeat transcripts. LL2R repeat-containing RNA also forms non-diffusible RNP-aggregates enriched by splicing factors that could serve as a site of post-transcriptional splicing. Our findings reveal how tandem repeat-derived RNAs drive the assembly of oocyte nuclear condensates, providing a functional template for understanding the formation of closely related nuclear speckles and nuclear stress bodies across eukaryotes.

## Introduction

Recent studies have shown that membraneless cellular compartments, both in the nucleus and in the cytoplasm, assemble through liquid-liquid phase separation (LLPS). LLPS is launched by local concentration of proteins containing internal disorder regions (IDR) (Banani et al. 2017; Uversky 2017; Wang et al. 2021; Yamazaki et al. 2022; Subedi et al. 2025). In the case of ribonucleoprotein (RNP)-rich membraneless cellular compartments, such as nuclear domains and cytoplasmic RNP granules, LLPS is facilitated and specified by RNA that binds RNA-binding proteins (RBPs) containing IDRs (Lin et al. 2015; Zhang et al. 2015; Langdon et al. 2018; Strom and Brangwynne 2019; Riback et al. 2020; Hirose et al. 2023). Among the many RNA components of different transient or stable nuclear RNP compartments, there are essential scaffold RNA species called architectural RNAs (arcRNAs), which are mostly long non-coding RNAs (ncRNAs) (Caudron-Herger and Rippe 2012; Chujo et al. 2016; Yamazaki et al. 2019; Yamamoto et al. 2020; Mattick et al. 2023; Hirose et al. 2025). To date, an architectural role in nuclear domain assembly has been experimentally demonstrated for a few long non-coding RNAs (lncRNAs): NEAT1_2 RNA that organises paraspeckles in mammalian cells (Clemson et al. 2009; Sasaki et al. 2009; Yamazaki et al. 2018; Takakuwa et al. 2023), transcripts of human satellite III in nuclear stress bodies (Valgardsdottir et al. 2005; Ninomiya et al. 2023), transcripts of ribosomal intergenic spacer in the nucleolar detention centre or amyloid body (Audas et al. 2012), Alu repeat-containing RNA in the nucleolus (Caudron-Herger et al. 2015), pyrimidine-rich noncoding transcript (PNCTR) in the perinucleolar compartment (Yap et al. 2018), hsr-ω-n RNA in Drosophila omega-speckles (Prasanth et al. 2000), meiRNA (Yamashita et al. 1998; Shichino et al. 2014) and mamRNA (Andric et al. 2021) in the Mei2/Mmi1 nuclear RNP foci in fission yeast, Gomafu/MIAT/Rncr2 in the Gomafu nuclear domain in a subset of neurons (Ip et al. 2016), and small nucleolar RNA (snoRNA) capped and polyadenylated lncRNAs (SPAs) at the Prader-Willi syndrome locus (Wu et al. 2016). Given that most characterized arcRNAs have non-conserved primary sequences and may have tissue-specific expression patterns, the expected list of nuclear arcRNAs is likely to be much longer and includes thousands of potential candidates (Quinodoz et al. 2021; Quinodoz and Guttman 2022; Yamazaki et al. 2022; Zeng et al. 2023; Hirose et al. 2025). However, to date, the range of characterised arcRNAs is very limited. Therefore, the search for novel spatially enriched ncRNAs that seed nuclear domain formation and the study of their role in the regulation of dynamic nuclear processes are of particular importance.

Growing oocyte nucleus in animals with hypertranscriptional type of oogenesis (fishes, amphibians, reptiles, birds and several invertebrates) is a promising model object for identifying new arcRNAs and studying mechanisms of nuclear domain formation (Krasikova et al. 2012; Krasikova and Kulikova 2019b). A major advantage of this model is the giant size of the nuclei and the transformation of the chromosomes into the ’lampbrush’ configuration, which is characterised by the appearance of thousands of transcription units that loop out laterally from the chromomere axis of the chromosome (Gaginskaya et al. 2009; Gall 2014; Morgan 2018; Krasikova et al. 2023). Chromosomes in a lampbrush form are characterized by intensive transcription of a variety of coding and non-coding DNA sequences, including tandemly repetitive DNA (Gardner et al. 2012; Trofimova and Krasikova 2016; Krasikova et al. 2024). Lampbrush chromosomes provide exceptional opportunities to map loci of nuclear domain formation with very high cytogenetic resolution, to analyse their ongoing transcription and nascent RNA processing at the cytological level, and to study their role in the nucleation of different loci-associated nuclear domains (Morgan 2002; Zlotina et al. 2016; Krasikova and Kulikova 2019b).

In addition to normal transcription loops bearing nascent gene transcripts (Krasikova et al. 2024), lampbrush chromosomes carry a number of complex loops, also called special or marker loops, with different morphology of their RNP-matrix (Callan 1986; Angelier et al. 1990; Morgan 2002; Krasikova and Kulikova 2019b). A prominent example of such loops are the so-called ‘lumpy loops’ that form on the long arm of lamprush chromosome 2 in chicken growing oocytes (Chelysheva et al. 1990). The dense RNP-matrix of ‘lumpy loops’ is strongly stained with Coomassie, indicating protein accumulation. Immunostaining of the whole nuclei isolated from chicken oocytes revealed that such complex loops belong to loci-specific RNP domains (Krasikova et al. 2012). Most of these nuclear domains concentrate splicing factors (Krasikova et al. 2004; Krasikova et al. 2012), which contain a low complexity IDR - arginine-serine (RS) rich domains that promote phase separation.

Here, we aimed to identify the non-coding RNAs responsible for scaffolding the splicing factor-enriched RNP domain associated with chicken lampbrush chromosome 2 and to establish the mechanism of its formation. We identified a novel single locus tandem repeat whose stable transcription from a retrotransposon promoter leads to the appearance of nuclear RNP condensates concentrating splicing factors. Our results indicate that a variety of tandem repeat-containing long non-coding RNAs facilitate nucleation of a diversity of membraneless nuclear compartments that recruit multiple sets of RNP complexes to defined genomic loci.

## Methods

### Bioinformatic analysis

The tandem repeat was identified by analysing the sequences bordering the gap between genomic contigs in the chicken genome assembly galGal3 (GCA_000002315.1) using the Ensembl (https://www.ensembl.org) and NCBI (https://www.ncbi.nlm.nih.gov/genome) Genome Browsers (Krasikova et al. 2010). Regions of homology with the identified tandem repeat in the assembled sequences of the chicken genome were then localised using Nucleotide BLAST (https://blast.ncbi.nlm.nih.gov/).

Chicken oocyte nuclear RNA and methylation profiles along telomere-to-telomere (T2T) chicken genome assembly GGswu (GCA_024206055.2) were obtained from our previous studies of chicken lampbrush stage oocyte transcriptome (Krasikova et al. 2024) and methylome (Nurislamov et al. 2022). CpG islands were identified using GC-Profile 2.0 with CpG observed/expected ratio >0.8 (Lai and Gao 2022). Tandem repeats were identified using Tandem Repeats Finder with Maxperiod 2000 (Benson 1999). Chicken repetitive elements from the Repbase library were localized using Censor (Kohany et al. 2006). Splicing sites were predicted using NNSplice v0.9 with a minimum score for 5’ and 3’ splice sites of 0.88 (Reese et al. 1997) and Spliceator v2.1 with options ’-td 97 -ta 97’ (Scalzitti et al. 2021). Only splice sites predicted by both programs were considered. RNA coverage tracks, positions of repetitive elements, and predicted splicing sites were visualized using the Integrative Genomics Viewer (Robinson et al. 2011). Calculations and visualizations were done with R software, version 4.2.2.

IntaRNA program (Mann et al. 2017) and QGRS Mapper (Kikin et al. 2006) were used to predict regions of RNA interaction.

### Molecular cloning and sequencing

Whole blood samples were collected from chicken (Gallus gallus domesticus) females, a commercial hybrid (cross) ISA brown. The animal studies received an approval of the Ethics Committee for Animal Research of St. Petersburg State University (#131-03-2, 14.03.2016; #131–04-6, 25.03.2019). Total genomic DNA was extracted from whole blood cells according to standard protocol (Maniatis et al. 1982). The LL2R repeat was amplified from genomic DNA by polymerase chain reaction (PCR) under the standard conditions with specific primers (Syntol):

446F: 5’- CCTGCAAGTAATGCGGTGAGC-3’;

446R: 5’-AGACAGCGGCCAACACC-3’.

The primers were designed according to the consensus sequence of tandem repeats bordering the gap between two contigs NW_001471654.1 and NW_001471655.1, at position 148,408,940 - 148,409,040 bp on GGA2 in chicken genome assembly galGal3 (GCA_000002315.1) (Krasikova et al. 2010).

The PCR product was separated by 1% agarose gel electrophoresis. The bright band of a fragment size of ∼385 bp, corresponding to a single unit (copy) of the repeat, was eluted from a gel using the MinElute Gel Extraction kit (Qiagen) and ligated into the pCR®II and the pCR®2.1 plasmid vectors (Invitrogen, Life Technologies). The E. coli strain Top10 was transformed with the ligation mixture, after which recombinant clones were selected by the blue-white screening and by PCR with LL2R-specific/M13 primers. Plasmid DNA was extracted from overnight bacterial cultures of the selected clones using the Wizard Plus SV Minipreps DNA purification system (Promega).

Clones containing LL2R inserts were bidirectionally sequenced using the BigDye Terminator v3.1 Cycle Sequencing Kit (Applied Biosystems, Life Technologies). The sequencing reaction was carried out according to the manufacturer’s recommendations with insignificant modifications. The sequencing mixture comprised 100 ng of the plasmid vector, 1.6 pM of M13 primer, 1.5 µl of BigDye Sequencing Buffer, and 1 µl of Ready Reaction Premix, with the total volume being 10 µl. The sequencing data obtained with the 3130 Genetic Analyzer (Applied Biosystems, Life Technologies) were further processed and analyzed using the Sequencing Analysis software (Applied Biosystems) and the Geneious software (www.geneious.com).

### Lampbrush chromosome preparations

Lampbrush chromosomes were manually isolated from growing oocytes with a diameter of 0.5-2.0 mm under the standard procedure (Kropotova and Gaginskaia 1984; Solovei et al. 1992). Oocyte nuclei were isolated in 5:1 medium (83 mM KCl, 17 mM NaCl, 6.5 mM Na2HPO4, 3.5 mM KH2PO4, 1 mM MgCl2, 1 mM DTT, pH 7.2). Nuclei were individually transferred to glass chambers mounted on the slides. Nuclear envelopes were removed in hypotonic 1/4 medium (5:1 diluted four times with the addition of 0.1% formaldehyde) with thin tungsten needles. Oocytes and nuclei were manipulated under a stereomicroscope Leica S9D or M165C (Leica Microsystems). Particular emphasis was given to the preservation of RNA during the isolation procedure. Nuclear contents were then spread by centrifugation (3000 rpm, 30 min at +4C°) in a cytology centrifuge (Hettich Universal 320R) and fixed in 2% PFA in 1×PBS (1.47 mM KH2PO4, 4.29 mM Na2HPO4, 137 mM NaCl, 2.68 mM KCl) for 15-30 min. Then, slides with chromosomes were dehydrated in an ethanol series and air-dried before FISH experiments or kept in a fresh solution of 70% ethanol before immunostaining.

Chicken mitotic metaphase chromosome spreads were prepared from embryonic fibroblasts according to standard protocols. Cells were treated with 0.1 μg/ml colcemid (Biological Industries) for 5 hours and then fixed in a cold 3:1 ethanol:acetic acid solution. The fixed cell suspension was dropped onto hot slides heated in a water bath, and then air dried (Deng et al. 2003).

### Staining of lampbrush chromosomes with SYTOX green

To reveal both DNA and RNA, lampbrush chromosome preparations were mounted with antifade medium (65% glycerol, 20 mM Tris-HCl, pH 7.5, 0.2% DABCO) containing the nucleic acid-specific fluorescent dye SYTOX green (Molecular Probes) and 1 μg/ml DAPI.

### Atomic force microscopy (AFM)

Preparations of lampbrush chromosomes were fixed with 2% formaldehyde in phosphate-buffered saline (1×PBS). The surface of lampbrush chromosomes was scanned using an atomic-force microscope Integra Aura (NT-MTD) in semi-contact mode with silicone cantilever (NSG01 and NSG10) according to the procedure described earlier (Krasikova and Kulikova 2016). Image analysis and 3D reconstructions were performed using the software NOVA (NM-MTD).

### Low-voltage scanning electron microscopy (SEM)

Preparations of lampbrush chromosomes were fixed by a mixture of 2.5% glutaraldehyde and 2% formaldehyde in 1×PBS. Preparations without conductive coating were analyzed in the scanning electron microscope Zeiss Merlin at low voltage (0.1–0.4lllkV) regime using a detector of secondary electrons (In-Lens SE). Scanning settings and image acquisition parameters are described in (Kulikova et al. 2016). Measurements were performed using the tools of the SmartTiff (Zeiss) software.

### Antibodies used

Following primary antibodies were used for immunofluorescent staining of lampbursh chromosome preparations: mouse monoclonal antibody (mAb) against double-stranded DNA (ab27156, Abcam), mAb V22 against the phosphorylated C-terminal domain (CTD) of RNA polymerase II (kindly provided by U. Scheer and R. Hock), mAb H14 against phosphorylated serine 5 of CTD repeat YSPTSPS of RNA polymerase II (Abcam, ab24759), mAb 4H8 against phosphorylated serine 5 of CTD repeat YSPTSPS of RNA polymerase II (Abcam, ab5408), mAb K121 against 2,2,7-trimethylguanosine (TMG) cap of splicing small nuclear RNAs (snRNAs) (Santa Cruz Biotechnology, Inc.), mAb Y12 against symmetric dimethylarginine of Sm proteins of spliceosomal of snRNPs (Lerner et al. 1981), mAb against serine/arginine-rich splicing factor SC35 and/or additional proteins of the spliceosome (Abcam, ab11826), mAb SCf11 against hypophosphorylated SC35 protein (Bubulya et al. 2004), mAb 4D11 against heterogeneous nuclear RNP protein L (hnRNP L) (Pinol-Roma et al. 1989), mAb 3C2 against hnRNP K (Matunis et al. 1992), rabbit polyclonal antibody against PTB-associated splicing factor PSF/SFPQ (Abcam, ab45359), rabbit polyclonal antibodies against hyperacetylated histone H4 (H4Ac) (06-866, Millipore), rabbit polyclonal antibodies against histone H3 trimethylated at lysine 9 (H3K9me3) (8898, Abcam), and mAb against 5-methylcytosine (5mC) (ab51552, Abcam).

For detection of primary antibodies, corresponding secondary antibodies were used: Cy3-conjugated goat anti-rabbit IgG, Alexa-488-conjugated goat anti-mouse IgG (Jackson Immuno Research Laboratories) and Cy3-conjugated goat anti-mouse IgG and IgM (Molecular Probes).

### Immunofluorescent staining

Immunofluorescent staining of lampbrush chromosomes was carried out as previously described (Krasikova et al. 2004). Briefly, after gradual rehydration in 70%, 50%, 30% ethanol to 1×PBS with 0.02% Tween-20 (Sigma), preparations were blocked in a blocking solution (0.5% blocking reagent (Roche) in 1×PBS), and then, successively incubated with primary and secondary antibodies in recommended dilutions in the blocking solution.

Incubations lasted one hour at RT. After incubation with antibodies, the preparations were washed 3 times in 1×PBS with 0.02% Tween-20 (Sigma). After immunostaining, preparations were dehydrated in an ethanol series, air dried, and mounted with antifade solution containing 1 μg/ml DAPI (Sigma). As a positive control part of the preparations selected for immunostaining with antibodies SCf11 were pretreated with 2.5 U of calf intestine alkaline phosphatase (Fermentas) at 50°C for 1h. The reaction was stopped by brief washing in 50 mM EDTA in 0.3M NaCl. Preparations selected for immunostaining with anti-5mC were denatured in 2 M HCl for 45-90 min at RT.

### Immunostaining and SYTOX green staining of isolated oocyte nuclei

Intact and non-deformed nuclei were manually isolated from healthy oocytes of 0.7-2.0 mm in diameter as described in the lampbrush chromosome preparation procedure. Isolated nuclei were transferred to 1×PBS and fixed with 2% formaldehyde in 1×PBS, followed by an immunostaining procedure (Krasikova et al. 2012). Then, 1×PBS was used for antibody dilution and as a washing solution for the nuclei. To visualize chromosomes and RNA-rich structures, unfixed intact nuclei were placed into individual coverslip chambers with 0.07 μM nucleic acid-specific fluorescent dye SYTOX green (Molecular Probes) in 5:1 medium (as in lampbrush chromosome isolation procedure). All manipulations with oocytes and nuclei were conducted under a stereomicroscope (Leica MZ16).

### Nascent RNA visualization by microinjection of BrUTP into oocytes

Oocytes dissected from the ovary were microinjected with 23 nl of 20 mM BrUTP in 1×PBS using authomatic nanoliter injector Drummond Nanoject II (Drummond Scientific) (Deryusheva et al. 2007; Kulikova, Chervyakova, et al. 2016). Lampbrush chromosomes were prepared from injected oocytes after 1-3 hours of incubation at 15°C in the humid chamber. BrUTP incorporated into newly synthesized transcripts was immunodetected with mAb 2B1 against BrdU (Santa Cruz Biotechnology, Inc.) according to the immunostaining procedure described above.

### FISH probes

BAC-clones positioned in the region of bioinformatically predicted locus of lumpy loop 2 formation, WAG-22B02, WAG-13J24, WAG-107K17 on GGA2, were selected from the Wageningen chicken BAC-clone library (Crooijmans et al. 2000). BAC CH261-49F17 to COL25A1 gene on GGA4 was selected from the CHORI-261 chicken BAC-clone library (https://bacpacresources.org/chicken261.htm). BAC-clones were amplified and labeled with biotin-16-dUTP or digoxigenin-11-dUTP by degenerated oligonucleotide-primed PCR (DOP-PCR) with 6-MW primer (Telenius et al. 1992) or by nick-translation (Green and Sambrook 2020) with DNase I/DNA polymerase I mix (Thermo Fisher Scientific).

The LL2R repeat probe was amplified and labeled with biotin-16-dUTP or digoxigenin-11-dUTP by PCR with specific primers (see ‘Molecular cloning and sequencing’).

The 5’-UTR of the CR1-H element probe was amplified and labeled with biotin-16-dUTP by PCR with specific primers (Lumiprobe):

CRH2_F: 5’-ACTGACCACGACATACAGCC-3’

CRH2_R: 5’-AGCCGAAGACCTGAACACTG-3’.

Oligonucleotide 5’-end biotin-labeled dT(30) probe (Syntol) was used to detect polyadenylated transcripts.

Strand-specific locked nucleic acid (LNA)-enhanced oligonucleotide probes to LL2R (Syntol) were used to identify the transcribed strand of LL2R:

446-2s: 5’-[Cy3]-TG+GTA+AGG+TCAT+TTATGC+CC+AC-3’;

446-2a: 5’-[FAM]-G+TG+GGCATA+AAT+G+ACC+TTA+CCA-3’;

446-3s: 5’-[Cy3]-GT+GAT+GAAT+T+TGCT+GGT+GTT+GGC-3’;

446-3a: 5’-[FAM]-GC+CAA+CA+C+CAGCA+AATT+CAT+CAC-3’.

LNA-enhanced oligonucleotide probe to U6 snRNA

5’-C+ACG+AAT+TTG+CGT+GTC+ATC+CTT-3’ (Syntol) was labeled by digoxigenin at the 3’-end. Symbol Ȝ+” precedes LNA-modified nucleotides.

Oligonucleotide probes to LL2R used for super-resolution SIM (Lumiprobe):

904-2a: 5’-[Cy3]-GGGTGGGCATAAATGACCTTACCAGGAGA-3’; 904-3a: 5’-[Cy5]-GCCAACACCAGCAAATTCATCACTGTACCA-3’.

Strand-specific oligonucleotide probes to CNM repeat (Krasikova et al. 2006): CNMneg: 5’-[Bio]-AAATGGGGGATTTTCGAAGAGAAAACA-3’;

CNMpos: 5’-[Cy3]-TGTTTTCTCTTCGAAAATCCCCCATTT-3’.

### FISH procedure

LL2R mapping. To identify the LL2R locus by physical mapping, DNA/DNA and DNA/(DNA+RNA) FISH protocols were used (Zlotina and Krasikova 2026). BAC-clones and LL2R PCR-based probes were denatured simultaneously with chromosomes (mitotic metaphase or lampbrush chromosomes).

LL2R transcripts detection. To detect LL2R transcripts and poly(A)+RNA on preparations of lampbrush chromosomes, the DNA/RNA FISH-protocol of (Kulikova and Krasikova 2022) was used. Chromosomes were not denatured, and RNase treatment was omitted.

Ribonuclease treatment assay. To examine the structural characteristics of LL2R repeat-containing RNA, lampbrush chromosome preparations were pretreated with ribonucleases of various specificities before DNA/RNA FISH with the LL2R-probe. RNase cocktail RiboShredder RNase Blend 50 U/ml (Epicenter Biotechnologies), RNase A 50 μg/ml (ThermoFisher Scientific), RNase H 100 U/ml (Epicenter Biotechnologies), RNase III 100 U/ml (Epicenter Biotechnologies), and RNase R 100 U/ml (Epicenter Biotechnologies) were applied as previously described (Trofimova et al. 2014).

Hybridization mixture contained: 20–40 ng/μl of a probe with a 50-fold excess of salmon sperm DNA (for PCR-based probes) or yeast tRNA (for oligonucleotide probes), 50% formamide (for PCR-based probes) or 42% formamide (for oligonucleotide probes), 10% dextran sulfate, and 2×SSC. PCR-based probes and/or chromosomes were denatured together at 80°C for 5 min in the hybridization mixture. For RNA-FISH, PCR-based probes were denatured separately at 95°C for 10 min. Hybridization was performed overnight at 37°C for PCR-based probes or at RT for oligonucleotide probes. Preparations hybridized with PCR-based probes were washed three times in 0.2×SSC at 60°C and two times in 2×SSC at 45°C. Preparations hybridized with oligonucleotide probes were washed three times in 2×SSC at 37°C. Biotin and digoxygenin were detected by standard procedure using Cy3- and Alexa488-conjugated avidin or antibodies against digoxygenin (Jackson Immuno Research Laboratories), respectively. After detection, preparations were dried and mounted in an antifade solution containing DAPI (1 μg/ml).

LNA-FISH. LNA-enhanced oligonucleotide probes were diluted in a hybridization buffer (500 ng/μl E. coli tRNA, 50% formamide, 10% dextran sulfate, 1×Denhardt’s Solution (ThermoFisher Scientific), 4.3xSSC), to a final concentration of 80 nM and denatured at 80°C for 5 min, placed on ice, and then applied to preparations of lampbrush chromosomes.

Hybridization lasted 2 h at 53°C. After hybridization preparations were washed with three changes of 0.2×SSC at 53°C, dehydrated in ethanol series, air dried, and mounted in antifade solution containing DAPI (1 μg/ml).

Immunostaining followed by RNA-FISH. Immunostaining was performed as described above, except that the blocking step was omitted and incubation solutions did not contain blocking reagent to preserve RNA. Some preparations were treated with 0.1% Triton X-100 in 1×PBS for 20 min and washed with three changes of 1×PBS before immunostaining and FISH to increase the permeability of the RNP-matrix for PCR-based probes. Immediately after immunostaining and dehydratation preparations were subjected to RNA-FISH procedure as described above.

### Widefield fluorescence microscopy

Lampbrush chromosome preparations were browsed and imaged using a fluorescent microscope, Leica DM4000B (Leica Microsystems, GmbH), equipped with the appropriate set of filter cubes. Images were acquired with a monochrome CCD camera DFC350 FX (Leica Microsystems, GmbH).

### Confocal laser scanning microscopy

Spatially (3D) preserved nuclei after SYTOX green staining or immunostaining with antibodies were analyzed in a confocal laser-scanning microscope Leica TCS SP5 (Leica Microsystems) based on an inverted microscope DIM6000BP equipped with a 40× oil immersion HCX PL APO objective, diode (405 nm), argon (488 nm), and helium-neon (543 nm) lasers. Fluorescence signals were acquired sequentially for each optical section. 3D reconstructions of the scanned nuclei were built from Z-stacks using the LAS AF software (Leica Microsystems) as described previously (Maslova and Krasikova 2011).

### Super-resolution structured illumination microscopy (SIM)

To examine the ultrastructural organization and composition of lampbrush chromosome transcription loops, spatial structured illumination microscopy (3D-SIM) was performed with a 63×/1.4 Oil Plan-Apochromat objective of an Elyra 7 microscope system and the software ZENBlack (Carl Zeiss GmbH). Images were captured separately for each fluorochrome using the 642, 561, 488, and 405 nm laser lines for excitation and appropriate emission filters (Weisshart et al. 2016). Lampbrush chromosome preparations were subjected to immunostaining and FISH procedures as described above.

## Results

### Lumpy loops on chicken lampbrush chromosome 2 represent RNP-enriched nuclear condensates corresponding to a single transcription unit

After staining of isolated chicken oocyte nuclei or lampbrush chromosome preparations with nucleic acid-specific dye Sytox Green, so-called ‘lumpy loops’ on the long arm of chromosome 2 fluoresce brightly. They represent units of the RNP-matrix with a high local concentration of RNA (Figures 1a-d). Morphologically lumpy loops are long loops of ∼20 μm in length, which emerge from a single chromomere with a size of ∼1-2 μm, with a single transcription unit producing a dense RNP-matrix (Figures 1d, d’). There are two pairs of ‘lumpy loops’, each emerging from both sister chromatids on each homologue. Lumpy loops are variable in size depending on the individual and sometimes show allelic differences by being larger on one homologue. Notably, lumpy loops remain visible after lampbrush chromosome condensation when most normal transcription loops are already retracted (Figures 1e, e’).

**Figure 1.**
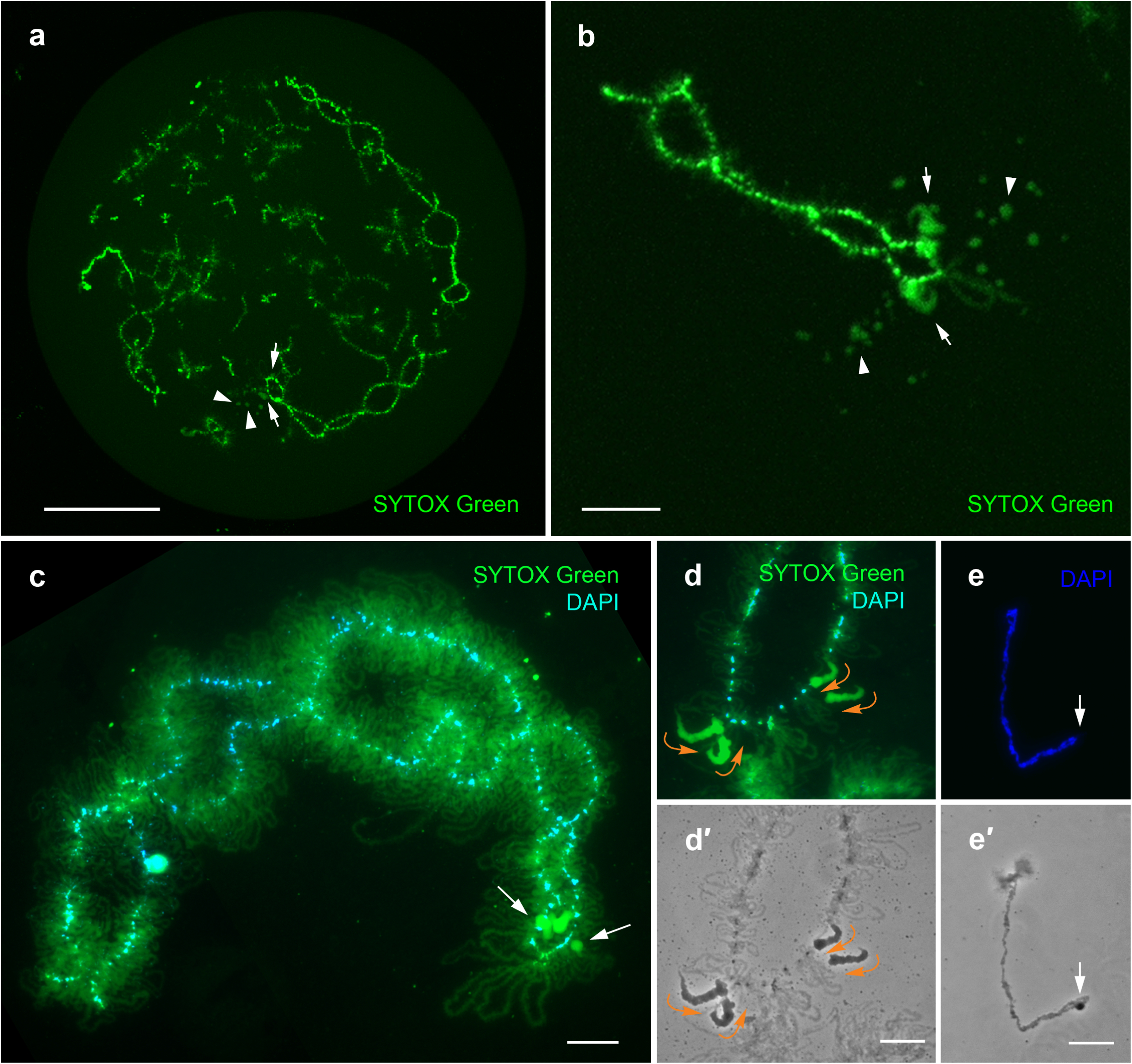
‘Lumpy loops’ on chicken lampbrush chromosome 2 represent RNP-enriched nuclear condensates corresponding to a single transcription unit. **a, b** – Maximal projections of isolated chicken oocyte nuclei stained with SYTOX green (green). Arrows indicate ‘lumpy loops’ on lampbrush chromosome 2, arrowheads indicate RNP-patches budding from the ‘lumpy loops’. Scale bars = 50 μm (a), 10 μm (b). **c, d** – Lampbrush chromosome 2 (c) and a fragment of lampbrush chromosome 2 with extended ‘lumpy loops’ (**d**) stained with SYTOX green (green) and DAPI (blue); **d**’ – corresponding phase contrast image. Arrows indicate ‘lumpy loops’, and orange curved arrows indicate the direction of transcription along ‘lumpy loops’. **e** – Condensed chromosome 2 at the end of the lampbrush stage. The arrow indicates a ‘lumpy loop’; **e**’ – corresponding phase contrast image. Scale bars = 10 μm.

Scanning of the chromosome 2 surface by both low-voltage scanning electron microscopy and atomic force microscopy revealed a significant volume and distinctive ultrastructural appearance of ‘lumpy loops’ dense fibrillar RNP-matrix in comparison to normal transcription loops (Figure 2; Supplementary Figure S1). ‘Lumpy loops’ start with a normal transcription unit that then changes the morphology of its RNP-matrix, producing longer RNP fibrils with a denser package. Despite a massive RNP-matrix, as opposed to the thinner RNP-matrix of normal lateral loops, these unusual transcription loops still form closed loops without flanks separation. Spherical budding RNP-aggregates of sizes up to 3 μm often arose from the ‘lumpy loop’ RNP-matrix (Figure 1a, b; Figure 2c, d).

**Figure 2.**
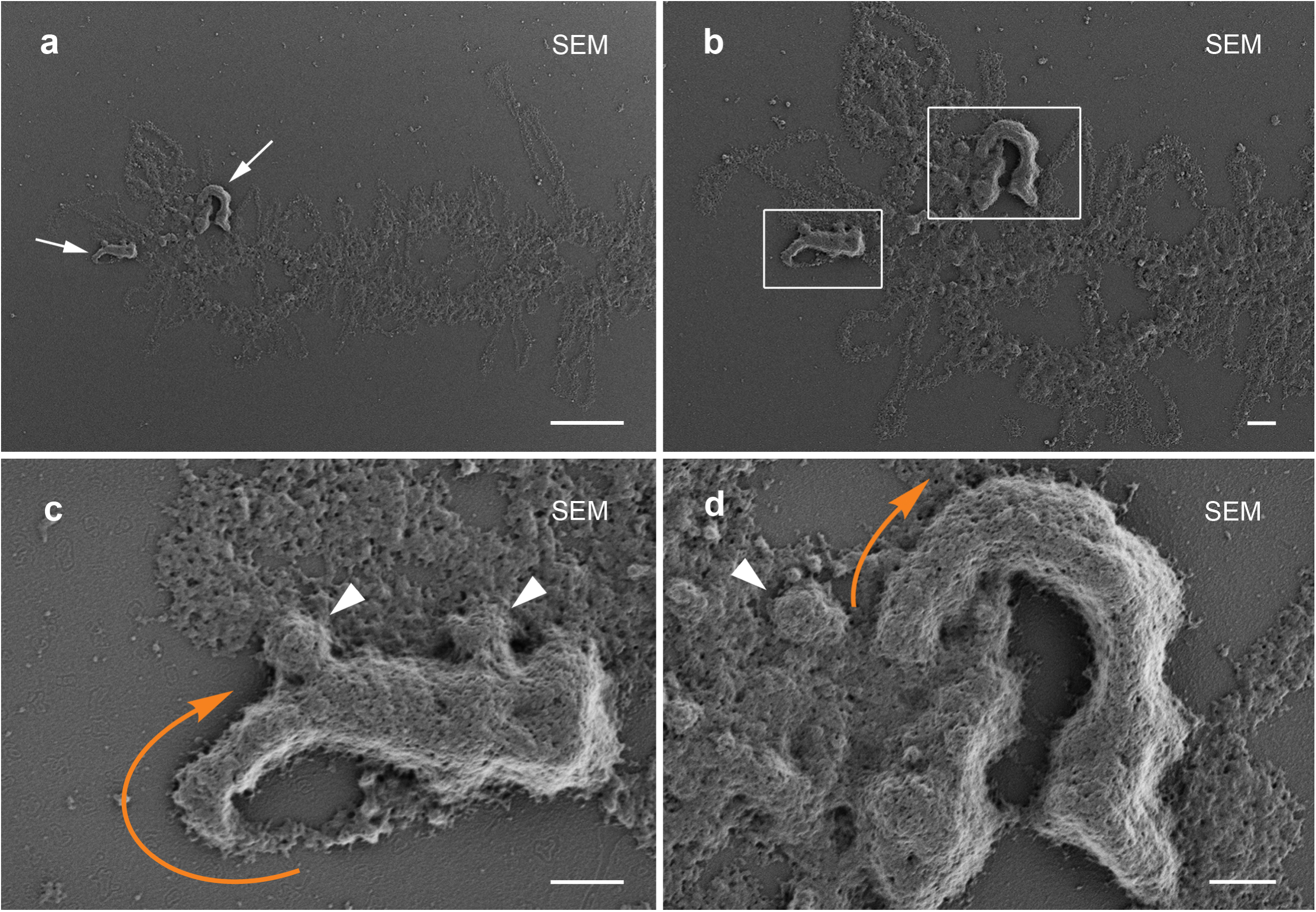
‘Lumpy loops’ on lampbrush chromosome 2 have a distinctive appearance in low-voltage SEM. **a, b** – Fragments of chicken lampbrush chromosome 2 visualized by low-voltage scanning electron microscopy (SEM); **c, d** – Enlargement of ‘lumpy loops’ framed in **b**. Arrows indicate ‘lumpy loops’, arrowheads indicate RNP-patches budding from ‘lumpy loops’, and orange curved arrows indicate the direction of transcription within ‘lumpy loops’. Scale bars = 10 μm (**a**), 2 μm (**b**), 1 μm (**d, c**).

### The genomic locus of ‘lumpy loop’ formation corresponds to a cluster of LL2R tandem repeats

The chicken chromosome 2 (GGA2)-specific painting probe labels brightly hundreds of normal lateral loops due to hybridization with nascent RNA, but labels only moderately the RNP-matrix of ‘lumpy loops’ because of pre-annealing with Cot-1 DNA containing the repetitive genomic fraction (Derjusheva et al. 2003). Thus, the RNA-content of the ‘lumpy loop’ transcription unit was apparently represented by repeat-enriched transcripts.

To identify nascent ‘lumpy loop’ transcripts, we first determined a genomic locus of their formation. At the first step, we performed DNA+RNA FISH on lampbrush chromosome 2 with BAC-clones WAG-22B02 and WAG-107K17 that contain sequences located on the long arm of GGA2. Both BAC-clones were mapped around the locus of ‘lumpy loop’ formation on chromosome 2 (Figures 3a, a’, f, f’). Next, we selected and hybridized another BAC-clone, WAG-13J24, closer to the loci of ‘lumpy loop’ formation. BAC-clone WAG-13J24 containing genomic sequences of the lncRNA gene ENSGALG00000047850 and the protein-coding gene ZFAT1 was mapped at the very beginning of the thin end of ‘lumpy loops’ (Figures 3a, a’, g). As a result, we determined that the ‘lumpy loop’ on lampbrush chromosome 2 corresponds to a gap in the assembled genomic sequences of chromosome 2 in the chicken genome version galGal3 (GCA_000002315.1) (Figures 3a-a’’).

**Figure 3.**
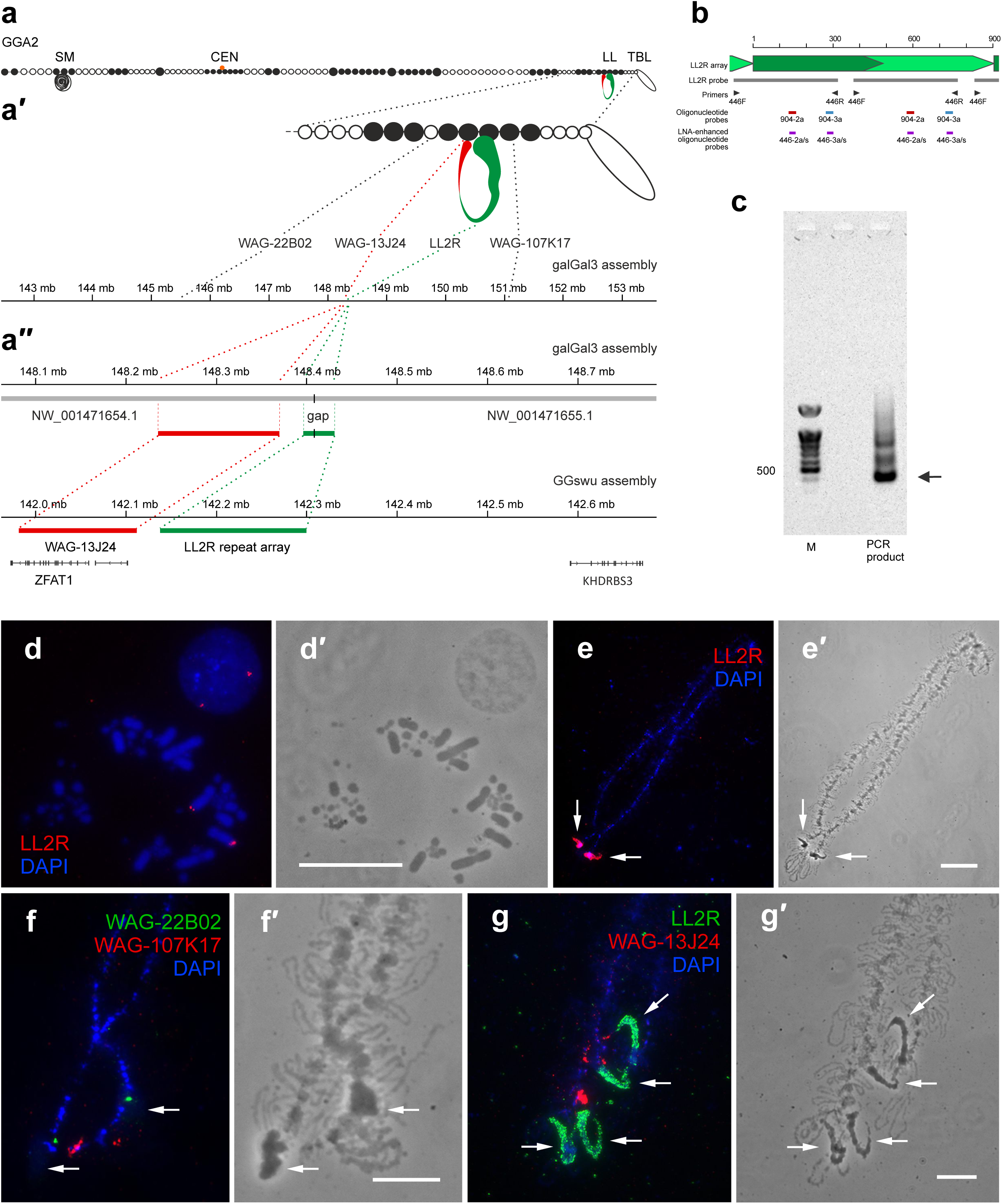
The genomic locus of ‘lumpy loops’ formation corresponds to a cluster of LL2R tandem repeats. **a - a**’’ – Correspondence of the locus of ‘lumpy loop’ formation to the cluster of tandem repeats. **a** – Cytological map of lampbrush chromosome 2, **a**’ – enlarged fragment of the cytological map with the positions of the LL2R tandem repeat cluster (148.4-148.43 Mb) and mapped BAC-clones (WAG-22B02 (145.44-145.55 Mb), WAG-13J24 (148.24-148.37 Mb), WAG-107K17 (151.1-151.2 Mb)) in the chicken genome assembly galGal3 (GCA_000002315.1). a’’ – Enlarged view of the region containing the LL2R cluster in chicken genome assemblies galGal3 and GGswu (GCA_024206055.2). In the galGal3 genome assembly, the LL2R repeat array is represented by a gap between contigs NW_001471654.1 and NW_001471655.1, surrounded by several copies of LL2R tandem repeats. **b** – The scheme illustrates positions of oligonucleotide and PCR-based probes to LL2R repeat relative to LL2R array monomers; PCR-based LL2R probe produced with primers 446F and 446R and corresponding fragment of LL2R monomer cloned from chicken genomic DNA (GenBank: KT952327, KT952328, KT952329). **c** – Detection of the LL2R sequence in the chicken genomic DNA using PCR with primers 446F and 446R; PCR products resolved in 1% agarose gel; lane M designates the size marker. **d** – DNA/DNA FISH with the LL2R-probe (red) on somatic metaphase chromosomes and in an interphase nucleus. A single cluster of LL2R repeat localises on the long arm of chromosome 2. **e** – DNA+RNA FISH on lampbrush chromosome 2 with the LL2R-probe (red). **f** – DNA+RNA FISH on lampbrush chromosome 2 with BAC-clones WAG-22B02 (green) and WAG-107K17 (red); **g** – DNA+RNA FISH on lampbrush chromosome 2 with BAC-clone WAG-13J24 (red) and the LL2R-probe (green). The LL2R repeat is located within ‘lumpy loops’ (arrows) on lampbrush chromosome 2. **d - g** – Chromosomes are counterstained with DAPI (blue). **d**’ - **g**’ – Corresponding phase contrast images. Scale bars = 10 μm (**d’, f’, g’**), 20 μm (**e**’).

Visual analysis of sequences bordering this gap allowed us to identify a novel tandem repeat with a monomer length of about 446 bp (Figure 3b). Next, we amplified this repeat from chicken genomic DNA by PCR with specific primers (Figure 3c). PCR fragments of ∼385 bp comprising a single unit of the repeat were cloned into plasmid vectors and sequenced (GenBank: KT952327.1, KT952328.1, KT952329.1). Probes for FISH-mapping were generated by PCR with the same primers (Figure 3b). FISH on somatic metaphase chromosomes revealed a single cluster of the identified repeat on the long arm of chromosome 2 (Figures 3d, d’). By DNA+RNA FISH on lampbrush chromosome preparations, we found that the identified tandem repeat maps precisely to the ‘lumpy loops’ on lampbrush chromosome 2 (Figure 3e, e’, g, g’). Thus, we named the repeat ‘Lumpy Loop 2 Repeat’ (LL2R).

Within the T2T chicken genome assembly (GGswu), the LL2R repeat array has 360 copies of ∼446 bp repeats, with an overall length of the array of ∼160 kb and the intra-array similarity of monomers of ∼86% (Figure 3a’’). In the genomic sequences of the LL2R repeat array, the Tandem Repeats Finder identified two types of repeats: the most prevalent TR904 monomer, consisting of two highly similar (88% identity) tandem repeats of ∼446 bp, and the TR9021 monomer, consisting of two less similar (56% identity) tandem repeats of ∼446 bp. Each TR904 monomer also contains one TTAGGG motif.

### Orthologous chromosomes in Galliformes species comprise LL2R-like repeats

The LL2R repeat is widespread in the order Galliformes, since using BLAST search of the NCBI Nucleotide collection (nr/nt) database, we were able to identify LL2R-like sequences in turkey (Meleagris gallopavo) (Supplementary Figure S2a), Japanese quail (Coturnix japonica), greater sage-grouse (Centrocercus urophasianus), common pheasant (Phasianus colchicus), helmeted guinea fowl (Numida meleagris), and western capercaillie (Tetrao urogallus) genomes. For instance, the turkey genome has a similar locus (between the ZFAT1 and KHDRBS3 genes) on the orthologous chromosome 3, which contains the tandem repeat TR1782 similar to the LL2R TR904 monomer (77.3% identity). The length of the TR1782 monomer is 1782 bp. The intra-array similarity of TR1782 monomers is ∼99%. The length of the array on turkey chromosome 3 is ∼10 kb (about five copies within the TR1782 array) (Supplementary Figure S2b). Interestingly, turkey lampbrush chromosome 3 does not bear any loops of special morphology within this locus (Myakoshina and Rodionov 1994); and Supplementary Figures S2c, c’) that can be explained by the low number of LL2R-like repeat copies. In the other representatives of Galliformes, the genomic region corresponding to the LL2R repeat array is represented by gaps of unknown size bordered by several copies of LL2R-like repeats, similar to other missing DNA in the reference assemblies of avian genomes (Peona et al. 2018).

### Nascent transcripts of the LL2R repeat nucleate the formation of lumpy loops

By RNA-FISH, we demonstrate that transcripts of LL2R repeats localize exactly in the ‘lumpy loops’ on lampbrush chromosome 2 (Figure 4a). The pretreatment of the chromosomes with the Riboschreder RNase cocktail or with RNAse A (which digests single-stranded RNA) eliminated LL2R RNA from the lumpy loops (Figure 4b, c). This result indicates that the LL2R tandem repeat is transcribed during oogenesis, and that transcripts take part in the formation of RNP-enriched nuclear domains (‘lumpy loops’) (Figure 4a).

**Figure 4.**
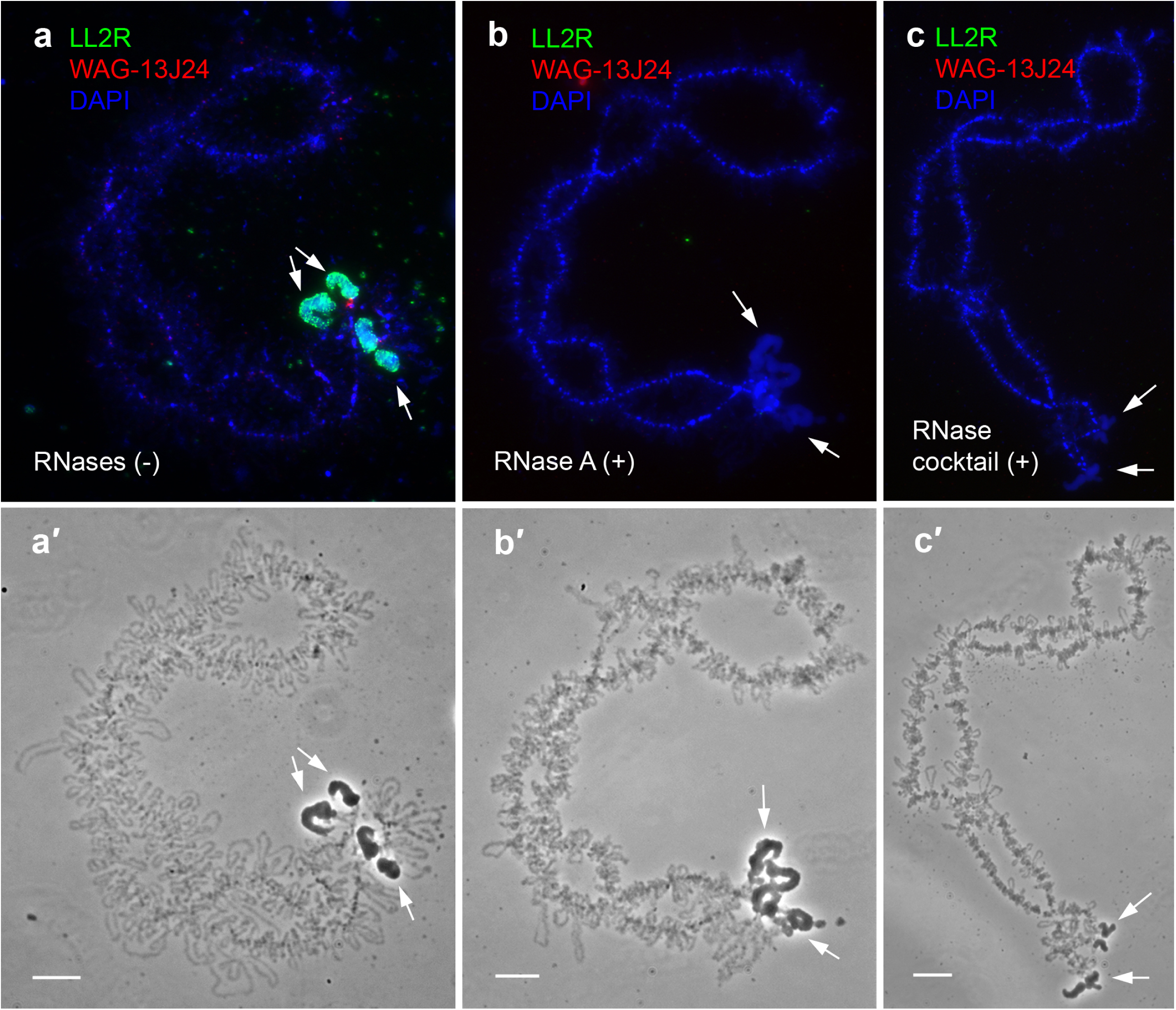
Transcripts of LL2R repeats are involved in the formation of ‘lumpy loops’. RNA-FISH with the LL2R probe (green) and BAC-clone WAG-13J24 (red) on chicken lampbrush chromosome 2; **a** – without any RNAse treatment; **b** – after treatment with RNAse A; **c** – after treatment with RNase cocktail (Riboschreder). Transcripts of LL2R repeats localize within ‘lumpy loops’ (arrows) (**a**) and can be eliminated by RNAse A (**b**). The effect of treatments with RNAses H, III, and R is shown in Supplementary Figure S3. Chromosomes are counterstained with DAPI (blue). **a**’- **c**’– corresponding phase contrast images. Scale bars = 10 μm.

Additional ribonuclease treatment assays revealed that LL2R transcripts are sustained after treatment with RNase H (that digests RNA from DNA/RNA hybrids), RNase III (that cleaves double-stranded RNA), or RNase R (that digests linear RNA from the 3’ end) (Supplementary Figure S3). We conclude that within ‘lumpy loops’, LL2R tandem repeat-containing RNAs are predominantly in a single-stranded form of nascent transcripts without free 3’ ends.

Moreover, the vast majority of all RNA at the LL2R locus is nascent RNA since no poly(A)-RNA is stored within the RNP-matrix of ‘lumpy loops’ according to RNA-FISH with an oligo-dT(30) probe (Supplementary Figures S4a-a’’) in contrast to poly(A)-rich RNP-aggregates at terminal regions of lampbrush chromosomes ((Kulikova, Chervyakova, et al. 2016) and Supplementary Figure S4b-b’’).

Next, we analysed potential RNA-RNA interactions between individual transcripts of the LL2R repeat. A partly complementary region was revealed between each copy of LL2R repeat RNA (TR904 consensus) (Supplementary Figure S5a). We also predicted the potential to form G-quadruplexes by LL2R repeat-containing RNA (Supplementary Figure S5b).

### The LL2R repeat array is transcribed on one strand from the LINE retrotransposon promoter

To identify the transcribed strand of the LL2R repeat array and the promoter region of this transcription unit, we analysed chicken lampbrush stage oocyte nuclear and cytoplasmic total stranded RNA-seq data aligned to the T2T chicken genome assembly (Krasikova et al. 2024). We established that the LL2R tandem repeat array is transcribed on one strand in a direction towards the telomere of the GGA2 q-arm (Figure 5a). Neighboring ENSGALG00000047850 and ZFAT1 genes are transcribed in the orientation opposite to the LL2R repeat array. RNA-FISH on lampbrush chromosome 2 with strand-specific LNA-enhanced oligonucleotide probes revealed only U-rich LL2R repeat repeat-containing transcripts (Figures 5d, d’, e, e’), thus confirming the stranded RNA-seq data.

**Figure 5.**
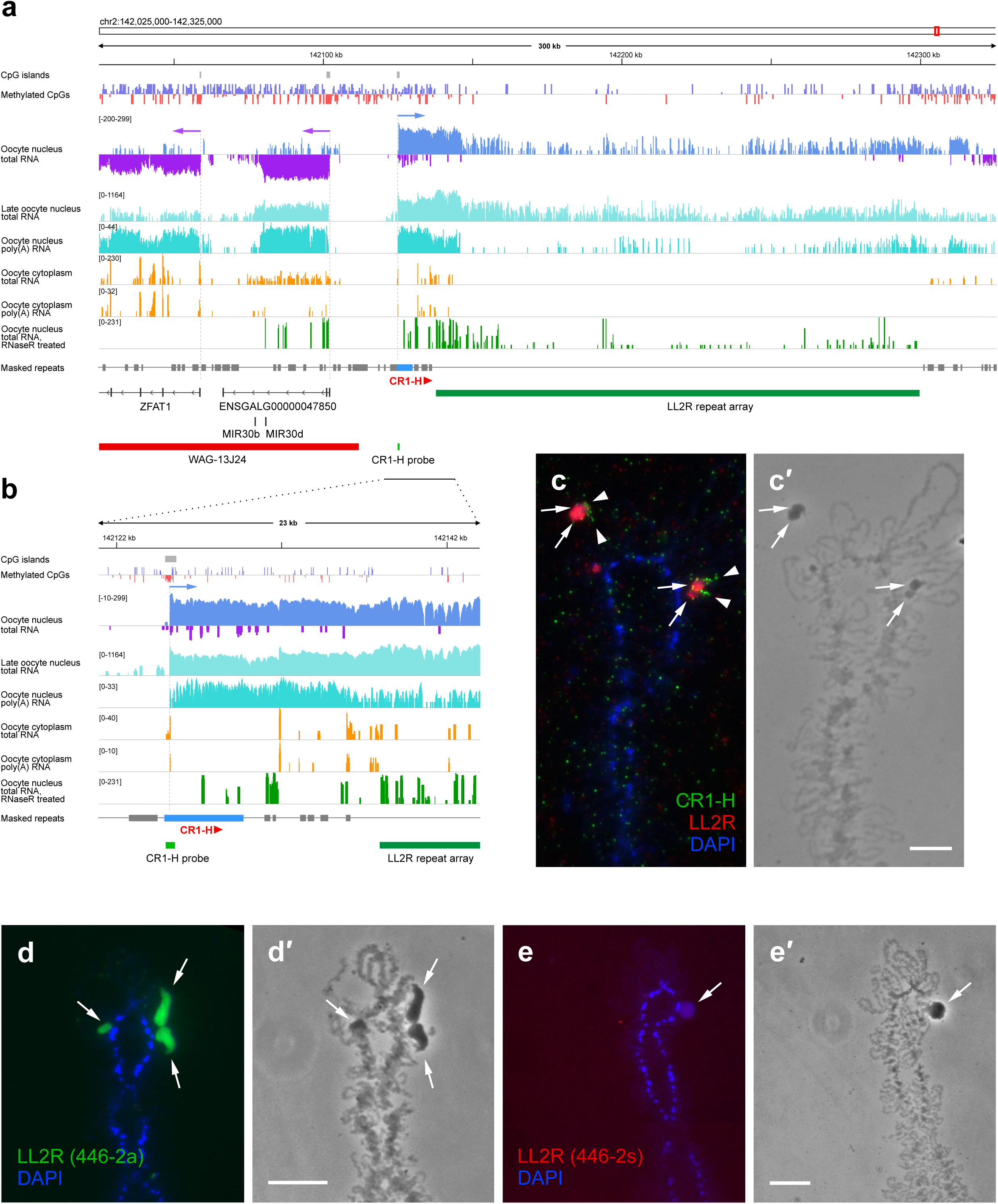
The LL2R repeat is transcribed on one strand from the CR1-H retrotransposon promoter. **a** – Organisation of transcription units in the vicinity of the LL2R repeat array. Overview of the GGA2 genomic region containing the LL2R repeat array and a fragment of BAC-clone WAG-13J24. Coverage tracks of total and poly (A) RNA from the nuclei and cytoplasm of lampbrush stage oocyte, as well as total RNA from the nuclei of post-lampbrush stage oocyte and total RNA from the nuclei of lampbrush stage oocyte after RNase R treatment, are visualized by the IGV browser and presented on a logarithmic scale. For total RNA from the lampbrush stage oocyte nuclei coverage track visualizes strand-specific read counts. The Methylated CpGs track visualizes methylome data of the single chicken diplotene oocyte nuclei (blue – methylated, red - demethylated); positions of CpG islands are shown on the individual track. The Masked repeats track visualizes the positions of repeats from the Repbase library. Full-length CR1-H element (corresponding to positions 1-4799 of the consensus) is located at the beginning of the transcription unit that encompasses the LL2R repeat array. LncRNA gene ENSGALG00000047850 and protein-coding gene ZFAT1 are located in the region preceding the LL2R repeat containing transcription unit. These genes are transcribed in the orientation opposite to the LL2R repeat array. 5’-ends of these genes are CpG islands that are demethylated. Vertical dotted lines indicate the boundaries of transcription units. Positions of genes, LL2R repeat array, BAC-clone WAG-13J24 and PCR-based probe to the 5’-end of CR1-H (corresponding to positions 51-595 of the consensus) are shown on individual tracks. **b** – Enlarged region shows the beginning of the transcription unit that encompasses the LL2R repeat array. Orientation of the CR1-H element corresponds to the direction of LL2R transcription. The 5’-end of this CR1-H is a CpG island that is demethylated and coincides with the transcription start site. **c** – Localisation of transcripts containing the 5’-end of the CR1-H element at the beginning of the LL2R repeat containing transcription loop. RNA-FISH with probes to the 5’-end of CR1-H (green) and LL2R (red). Chromosomes are counterstained with DAPI (blue). c’ – corresponding phase contrast image. Arrows indicate ‘lumpy loops’ on lampbrush chromosome 2, arrowheads indicate signal from the probe to the 5’-end of CR1-H. Scale bar = 10 μm. **d, e** – Identification of the transcribed strand of the LL2R repeat with strand-specific LNA-enhanced oligonucleotide probes. RNA-FISH with 446-2a probe for detection of transcripts identical to the positive strand (green) (d) and 446-2s probe for detection of transcripts identical to the negative strand (red) (e), d’, e’ – corresponding phase contrast images. Arrows indicate ‘lumpy loops’ on lampbrush chromosome 2. Chromosomes are counterstained with DAPI (blue). Scale bars = 10 μm.

In the total nuclear RNA fraction from lampbrush stage oocytes, the level of LL2R transcripts is comparable to the level of transcripts of the neighboring ENSGALG00000047850 and ZFAT1 genes (Figure 5a). In the total nuclear RNA fraction from post-lampbrush stage oocytes, the level of LL2R transcripts is considerably higher. Transcripts of LL2R repeat are also present in the oocyte nucleus poly(A) RNA fraction, implying that released LL2R repeat-containing RNA could be polyadenylated. The appearance of reads along the LL2R repeat array in the RNase R-treated oocyte nuclear total RNA fraction (Figure 5a) is consistent with the RNA-FISH data (Supplementary Figure S3c) and indicates an enrichment of nascent LL2R repeat-containing RNA.

At the beginning of the transcription unit of the LL2R tandem repeat array, we found a full-length LINE retrotransposable element of the CR1-H subfamily, 5’UTR of which could serve as a promoter (Figure 5a). According to our total stranded oocyte nuclear RNA-seq data, transcription of the LL2R tandem repeat array is indeed initiated at the 5’UTR of CR1-H element (Figure 5a). The distance from the beginning of the transcriptional unit (CR1-H 5’UTR) to the beginning of the LL2R array is ∼13 kb. RNA FISH with a PCR-based probe to the 5’-end of CR1-H confirmed its localization and transcriptional activity at the beginning of the lumpy loop bearing the LL2R repeat array (Figures 5c, c’). Likewise, a nearly full-length CR1-H element precedes a short LL2R-like repeat array in the turkey genome (Supplementary Figure S2b).

The methylation profile for chicken lampbrush stage oocyte nuclei (Nurislamov et al. 2022) aligned to the T2T chicken genome assembly demonstrates that the 5’UTR of the full-length CR1-H element at the beginning of the LL2R repeat transcription unit is demethylated (Figure 5b), indicating its active promoter function in growing oocytes. 5’-ends of lncRNA gene ENSGALG00000047850 and protein-coding gene ZFAT1, located in the region preceding the LL2R repeat array, are also demethylated as expected (Figure 5a).

### The LL2R repeat transcriptional unit is covered by elongating RNA polymerases but demonstrates a low rate of nascent RNA synthesis

To explore the transcriptional activity of LL2R transcription loops, we first examined the distribution of the active form of RNA polymerase II and then followed the dynamics of incorporation into nascent transcripts of BrUTP injected into oocytes. Using antibodies H14, V22, and 4H8 against the hyperphosphorylated C-terminal domain of RNA polymerase II, followed by FISH with the LL2R probe, we revealed the hyperphosphorylated form of RNA polymerase II in a punctate pattern on the axes of LL2R transcription loops (Figures 6a-a’’’, b, b’). RNA polymerase II complexes were distributed more densely along the ’lumpy loop’ axes than on the axes of the other, normal transcription loops, as revealed by super-resolution 3D-SIM (Supplementary Figures S6a, b).

**Figure 6.**
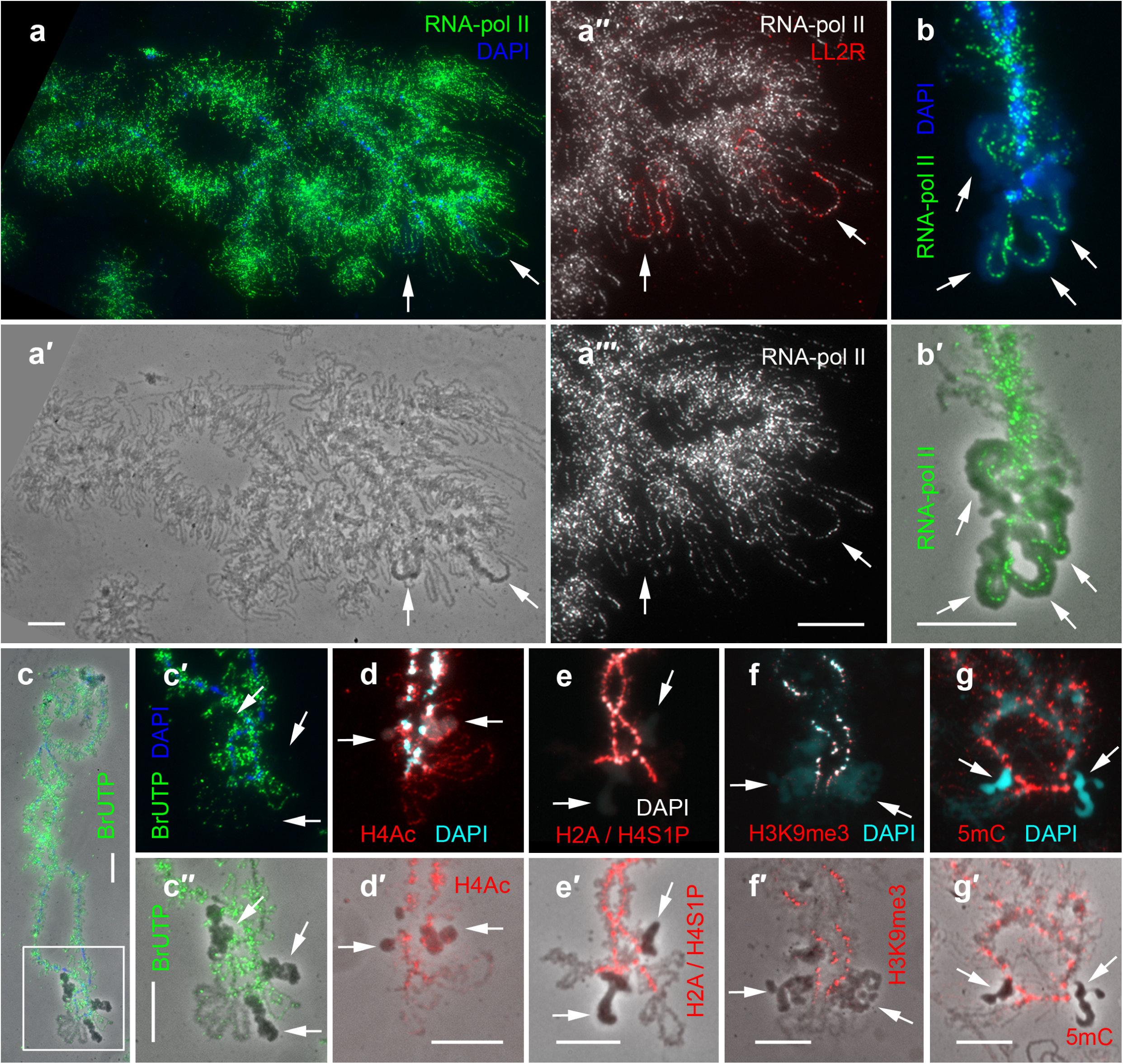
The LL2R transcriptional unit is a transcriptionally active region of chromatin with a low rate of nascent RNA synthesis. **a, b** – Immunodetection of hyperphosphorylated C-terminal domain of RNA-polymerase II with V22 antibodies; **a** – Full-sized lampbrush chromosome 2, V22 (green), DAPI (blue); **a**’ – corresponding phase contrast image; **a**’’, **a**’’’ – Fragment of lampbrush chromosome 2 shown in **a**, V22 (white) and DNA/DNA FISH with the LL2R probe (red). **b** – Fragment of lampbrush chromosome 2 after immunodetection of hyperphosphorylated C-terminal domain of RNA polymerase II (green), DAPI (blue); **b**’ – phase contrast image merged with immunofluorescent image (green). DNP axes of ‘lumpy loops’ (arrows) are strongly labeled. **с** – Immunodetection of BrUTP (green) incorporated into nascent transcript two hours after injection into the oocyte, DAPI (blue), **c**’ – enlarged fragment of **c**; **c**’’ – Phase contrast image merged with BrUTP fluorescence signal (green). ‘Lumpy loops’ (arrows) are devoid of labelling in contrast to normal lateral loops, which are labelled uniformly. **d** – Immunodetection of hyperacetylated histone H4 (red). **e** – Immunodetection of histones H2A and H4 phosphorylated at serine 1 (red). **f** – Immunodetection of histone H3 tri-methylated by lysine 9 (red). **g** – Immunodetection of 5-methyl cytosine (red). **d-g** – chromosomes are counterstained with DAPI (blue). **a’-g’** – Corresponding phase contrast images merged with immunofluorescence signals. ’Arrows indicate ‘lumpy loops’. Scale bars = 10 μm.

At the same time, LL2R transcription loops did not demonstrate any significant incorporation of BrUTP during 3h of incubation of injected oocytes, whereas normal transcription loops incorporated BrUTP into their nascent transcripts within the first 20 minutes after injection (Figures 6c-c’’). The distribution of RNA polymerase II, together with the dynamics of BrUTP incorporation suggest RNA polymerases stalling along the DNP-axes of ‘lumpy loops’ and a very low rate of nascent RNA synthesis. Thus, we propose that transcripts of the LL2R repeat tend to be retained at their transcription unit and are poorly released from the transcription loop.

### Epigenetic status of chromatin axes of LL2R repeat transcription loops

Despite the unusually low rate of RNA synthesis, loops formed by the LL2R-repeat transcription unit are not enriched with markers of repressed chromatin such as 5-methyl cytosine and histone H3 tri-methylated by lysine 9, which instead accumulate within chromomeres (Figures 6f, f’, g, g’). The LL2R repeat transcription loops also do not accumulate histones H2A and H4 phosphorylated at serine 1 (Figures 6e, e’), which is a marker of chromatin condensation. In contrast, hyperacetylated histone H4 is present in LL2R transcription loops in a punctate pattern, similar to most normal transcription loops (Figures 6d, d’).

Contour length of the transcription unit occupied by the LL2R repeat array measured after DNA/DNA-FISH (for example, Figure 6 a’’) is up to 18 μm. If we take into account the overall length of the LL2R repeat array in the T2T chicken genome assembly (GGswu) (∼160 kb), then the compaction level of its transcription unit is ∼8.8 kb/μm. The estimated deoxyribonucleoprotein (DNP) package level for this genomic region without nucleosomes is 3 kb/μm, and with nucleosomes it is 20 kb/μm. Thus, we conclude that the DNP-axes of the LL2R repeat transcription unit is highly decondensed with only rare nucleosomes, as confirmed by super-resolution 3D-SIM of immunodetected double-stranded DNA (Supplementary Figures S6c, d).

### Nascent transcripts of the LL2R tandem repeat recruit splicing factors to locus-specific RNP condensates

The composition of the LL2R transcript containing RNPs in ‘lumpy loops’ is different from the composition of pre-mRNPs in normal transcription loops. Namely, hnRNP L, hnRNP K, and SFPQ, widely distributed in the RNP-matrix of normal transcription loops bearing nascent gene transcripts, are specifically absent from the RNP-matrix of the LL2R repeat transcription unit (Figures 7e-g). Thus, not all nascent RNA processing factors bind LL2R transcripts and associated RBPs co-transcriptionally.

**Figure 7.**
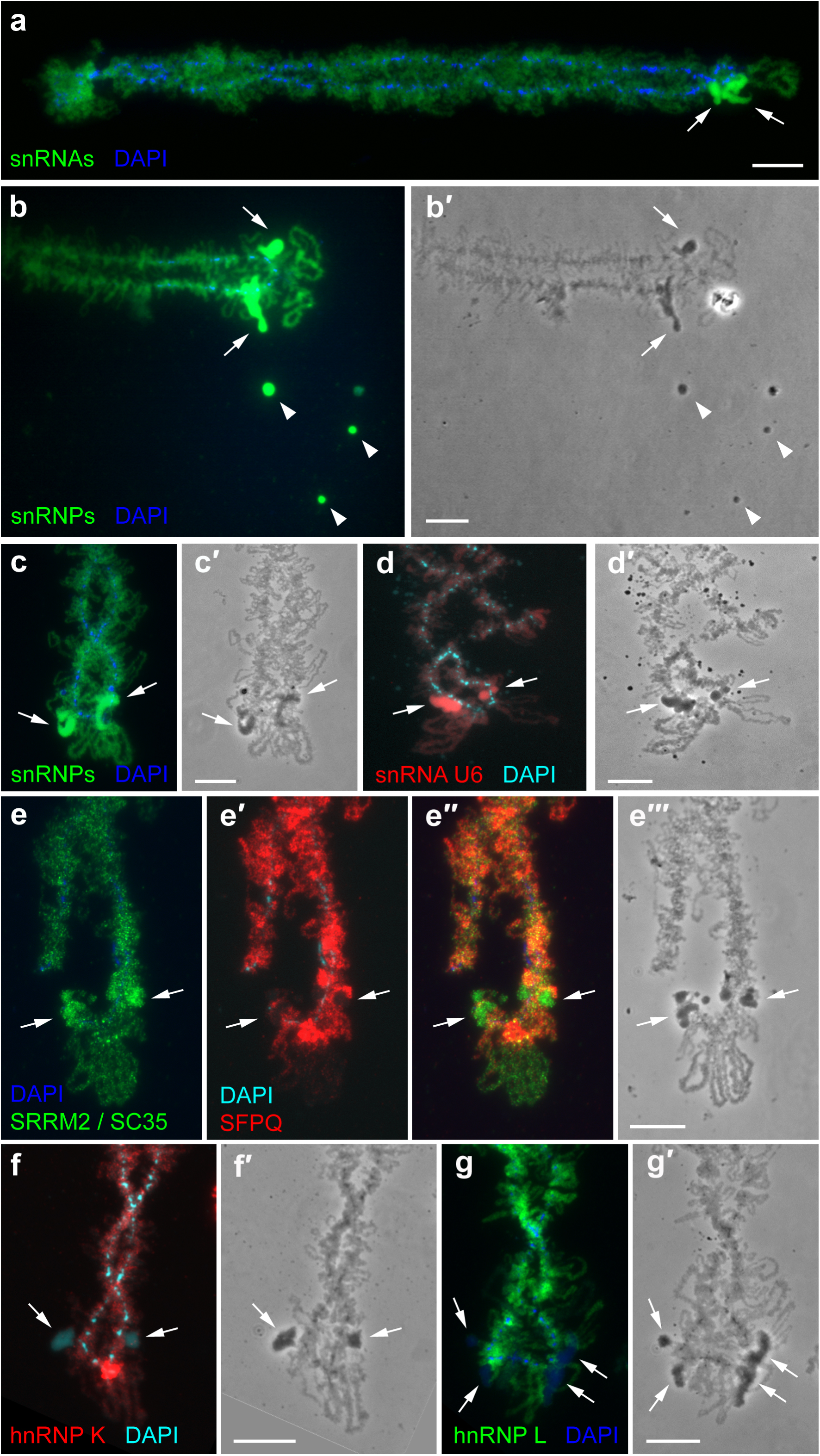
RNP composition of ‘lumpy loops’ bearing transcripts of the LL2R repeat. **a** – Immunodetection of TMG-cap of snRNAs with K121 antibodies; **b, c** – Immunodetection of Sm proteins of snRNPs with Y12 antibodies; **d** – RNA-FISH with LNA-probe to U6 snRNA; **e-e**’’’ – Double immunostaining with antibodies against splicing factors SRRM2 and SC35 (green) (**e**) and PSF/SFPQ (red) (**e’**); **e’’** – Merged image; **e**’’’ – Phase contrast image; **f** – immunodetection of hnRNP K (red); **g** – Immunodetection of hnRNP L (green). Chromosomes are counterstained with DAPI (blue). **b’- d’, f’, g’** – corresponding phase contrast images. ‘Lumpy loops’ (arrows) accumulate core splicing factors but lack hnRNP L, hnRNP K, and SFPQ. Arrowheads – RNP-aggregates enriched with splicing factors. Scale bars = 10 μm.

At the same time, LL2R transcripts are colocalized with high concentrations of mature snRNAs containing a tri-methylguanosine cap (including U6 snRNA detected by RNA FISH), core snRNP proteins containing glycine-arginine rich domain, serine/arginine repetitive matrix protein 2 (SRRM2) and SR phospho-protein SC35, which are required for splicing (Figure 7a-e). 3D-SIM confirmed the accumulation of splicing factors in the RNP-matrix of the LL2R repeat transcription units at the ultrastructural level (Supplementary Figure S7). Within ‘lumpy loops’, SC35 appears in its phosphorylated form as revealed by immunodetection of hypophosphorylated SC35 only after treatment with phosphatase (Supplementary Figure S8). These splicing factors are also generally distributed in normal transcription loops along their entire length. Additionally, 3D immunostaining at the intact oocyte nucleus demonstrated the accumulation of spliceosomal components within ‘lumpy loops’ on lampbrush chromosome 2 compared to nucleoplasm and normal transcription loops. Therefore, noncoding RNA of the LL2R repeat co-transcriptionally recruits RNA splicing factors, thus concentrating a specific set of RBPs within a locus-associated nuclear domain.

### High density of predicted donor splice sites in the LL2R tandem repeat-containing RNA

We then analyzed potential splice sites in the LL2R tandem repeat-containing RNA compared to pre-mRNA (Figures 8a, c). Immunolabeling of TMG cap of snRNAs followed by RNA-FISH with a BAC probe against the COL25A1 gene on lampbrush chromosome 4 demonstrated co-transcriptional splicing factors recruiting to nascent gene transcripts (Figures 8b, b’’). 3D-SIM revealed at least one thin-to-thick gradient of the RNP-matrix within the COL25A1 gene transcription loop, corresponding to a long intron. Nascent COL25A1 gene transcripts co-localized only partially with spliceosomal snRNAs within the RNP-matrix of the lateral loop (Figures 8b’, b’’). The COL25A1 gene body has comparable densities of predicted donor and acceptor splice sites (1-1.5 donor and acceptor splice sites per 2 kb length) (Figure 8a).

**Figure 8.**
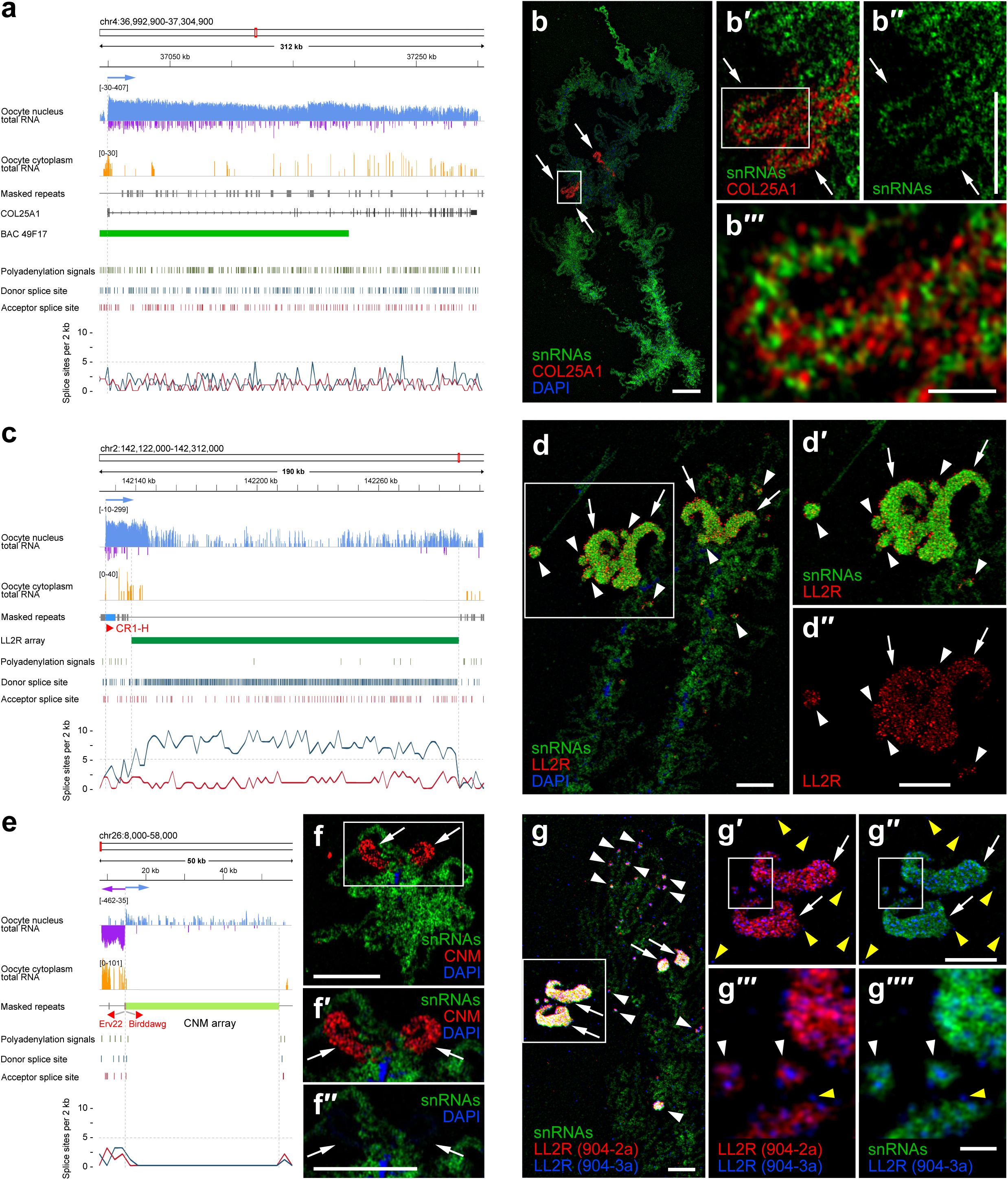
High density of donor splice sites in the LL2R tandem repeat containing RNA leads to concentration of splicing factors on ‘lumpy loops’ and in RNP-aggregates bearing transcripts of LL2R repeats. **a, c, e** – Different densities of predicted splice sites and polyadenylation signals across COL25A1 gene, LL2R repeat array and CNM repeat array. Overview of the GGA4 region containing the COL25A1 gene (**a**), overview of the GGA2 region containing the LL2R repeat array (**c**), overview of the subtelomere region of GGA26 containing the CNM repeat array (**e**). Coverage tracks of total RNA from the oocyte nucleus (strand-specific read counts) and cytoplasm (Krasikova et al., 2024) are visualized by the IGV browser and presented on a logarithmic scale. The Masked repeats track visualizes the positions of repeats from the Repbase library. Positions of the COL25A1 gene, LL2R repeat array and CNM repeat array are shown on individual tracks. The orientation of the selected CR1 and LTR-containing elements is indicated by the symbols ‘>’ for the positive and ‘<’ for the negative strand. The names of CR1 and LTR-containing elements whose position coincides with the boundaries of transcription units are shown in red. Vertical dotted lines indicate the boundaries of selected transcription units or tandem repeat arrays. Individual predicted donor and acceptor splice sites and polyadenylation signals are shown by vertical bars. The graphs below show the average (in a 2 kb window) density of predicted splice sites. Note the enrichment of the donor splice sites within the LL2R repeat array and the absence of the splice sites within the CNM repeat array. **b, d, f, g – 3D-SIM of COL25A1 gene, LL2R repeat array and CNM repeat array transcription loops on chicken lampbrush chromosomes.** Immunolabeling of TMG cap of snRNAs (green) followed by RNA-FISH with BAC probe CH261-49F17 to COL25A1 gene (red) on lampbrush chromosome 4 (**b**), PCR probe to the LL2R repeat (red) on lampbrush chromosome 2 (**d**), CNMneg probe to CNM repeat (red) on a microchromosome (**f**). Chromosomes are counterstained with DAPI (blue). Note the absence of spliceosomal snRNAs on the CNM repeat transcription loops and the enrichment of spliceosomal snRNAs on the LL2R repeat transcription loops. Immunolabeling of the TMG cap of snRNAs (green) followed by RNA-FISH with oligonucleotide probe 904-2a overlapping with the predicted donor splice site of the LL2R repeat (red) and oligonucleotide probe 904-3a overlapping with the region of the LL2R repeat devoid of splice sites (blue) (**g**). Transcription loops framed on **b, d, f, g** are enlarged in **b’, b’’, d’, d’’, f’, f’’, g’, g’**’. Areas framed on b’, g’, g’’ are enlarged in **b’’’, g’’’, g’’’’**. Arrows indicate transcription loops, arrowheads indicate RNP-aggregates, and yellow arrowheads indicate RNP-globules. Scale bars = 10 μm (**b**), 5 μm (**b’’, d, d’’, f, f’’, g, g’’**), 1 μm (**b’’’, g’’’’**).

Analysis of the LL2R tandem repeat-containing RNA also revealed potential splice sites. However, in contrast to protein-coding genes, the transcript of LL2R repeat array contains more than threefold excess of predicted donor splice sites compared to acceptor splice sites (1-1.5 acceptor and more than 5 donor splice sites per 2 kb length), and is depleted in polyadenylation signals (Figure 8c). Thus, we suggest that splicing of LL2R repeat-containing RNA can not be completed correctly during transcription due to an unbalanced number of donor and acceptor splice sites. At the same time, the CR1-H element is spliced out co-transcriptionally from nascent LL2R repeat-containing RNA as its transcript is not revealed within the RNP-matrix of the ‘lumpy loops’, being localized only at the beginning of the transcription unit (Figures 5c, c’). The LL2R-like repeat array on turkey chromosome 3, which does not produce lumpy loops at the lampbrush stage, has balanced densities of putative donor and acceptor splice sites (Supplementary Figure S2).

3D-SIM of lampbrush chromosome preparations after immunodetection of TMG cap of snRNAs followed by RNA-FISH with a probe to the LL2R repeat demonstrated partial co-localization of spliceosomal snRNAs and nascent transcripts of tandem LL2R repeat (Figures 8d-d’’).

In contrast, as it was shown by immuno-FISH on chicken lampbrush chromosome preparations, transcription loops of short 41-bp tandem repeat arrays lack spliceosomal snRNAs (Deryusheva et al. 2007). We found that both donor and acceptor splice sites are completely absent in a number of transcribed CNM repeat arrays in the chicken T2T genome assembly, for example, in the subtelomere region of microchromosome 26 (Figure 8e).

Accordingly, 3D-SIM demonstrated a complete absence of spliceosomal snRNAs on C-rich nascent transcripts of tandem CNM repeat on the lateral loops extending from centromere-associated chromomeres on several microchromosomes in a lampbrush form (Figures 8f, f’’). We conclude that the high concentration of splicing factors on lumpy loops transcribing the LL2R repeat array is due to the high density of donor splice sites in the LL2R tandem repeat-containing RNA.

### Non-diffusible RNP-aggregates are radially distributed around the site of LL2R repeat transcription

In many transcriptionally active nuclei, patches of non-diffusible RNP-aggregates not associated with chromosomes can be found in the nucleoplasm around ‘lumpy loops’ (Figures 1a, b; 2c, d). These derivatives accumulate large amounts of non-coding transcripts of the LL2R repeat (Figures 8d, d’’; Supplementary Figure S6a) with associated splicing factors such as snRNPs and SRRM2 (Figures 7 b, b’; 8d, d’; Supplementary Figure S7) but lack DNP-axes and RNA polymerase II. LL2R repeat-containing RNA within RNP-aggregates can be eliminated by treatment with RNase A but not by RNase H or RNase R treatment (Figure 4b; Supplementary Figure S3). Thus, LL2R repeat transcription units produce often a large amount of RNP-aggregates radially distributed around the site of transcription.

3D-SIM revealed DNA/RNA hybridization signals from the long PCR probe to the LL2R repeat mostly on the surface of ‘lumpy loops’ and RNP-aggregates (Supplementary Figures S7b’-d’). However, pretreatment of lampbrush chromosome preparations with 0.1% Triton X-100 before RNA-FISH allowed revealing hybridization signals within the inner part of ‘lumpy loops’ and RNP-aggregates (Supplementary Figure S7a’). The RNP-matrix of the ‘lumpy loops’ and RNP-aggregates radially distributed around ‘lumpy loops’ consists of RNP-fibers and RNP-globules comprising LL2R repeat-containing RNA.

Next, we performed immunodetection of TMG-cap of mature spliceosomal snRNAs followed by RNA FISH with two different oligonucleotide probes to the LL2R repeat on lampbrush chromosome preparations and analyzed their co-localization by 3D-SIM. Oligonucleotide probe 904-2a-Cy3, overlapping with predicted donor splice site and the region that can form RNA-RNA interactions (Supplementary Figure S5c), labels nascent LL2R repeat transcripts within ‘lumpy loops’ and in RNP-aggregates formed by RNP-fibrils that colocalize with spliceosomal snRNAs (Figures 8g, g’). At the same time, oligonucleotide probe 904-3a-Cy5, overlapping with the region devoid of splice sites (Supplementary Figure S5c), labels not only LL2R repeat-containing nascent RNP-fibrils but also small RNP-globules on the surface of ‘lumpy loops’, within RNP-aggregates, and free in the nucleoplasm (Figures 8g-g’’; Supplementary Figures S6c, c’, d). Small LL2R repeat containing RNP-globules are labeled only by the 904-3a-Cy5 oligonucleotide probe but not by the 904-2a-Cy3 probe (Figure 8g’). Moreover, small LL2R repeat-containing RNP-globules do not contain spliceosomal snRNAs (Figure 8g’’). We also noticed that small RNP-granules on the surface of ‘lumpy loops’ contain poly(A) RNA tail according to RNA-FISH with the oligo-dT(30) probe (Supplementary Figures S4a, a’). Thus, we conclude that partially processed LL2R repeat-containing transcripts form non-diffusible RNP-aggregates enriched by splicing factors that could serve as a site of post-transcriptional splicing. At the same time, completely processed polyadenylated LL2R repeat-containing RNAs diffuse freely in the nucleoplasm.

## Discussion

Here, using a combination of T2T genomic, transcriptomic, and cytomolecular approaches, we revealed the array of 445 bp tandem repeat LL2R on chicken chromosome 2, which is slowly transcribed as a single transcription unit from a retrotransposon promoter. By scanning electron and super-resolution microscopy for ultrastructural analysis, we demonstrate that nascent LL2R repeat transcripts are retained at their transcription sites, seeding the formation of a nuclear domain enriched with core splicing factors. The data indicate the role of tandem repeat-containing RNAs in the formation of oocyte nuclear condensates closely related to nuclear speckles or nuclear stress bodies in interphase nuclei of somatic cells.

### Transcription and processing of tandem repeat-containing RNA

The question of how transcription of tandemly repeated DNA is initiated and terminated remains open (Smurova and De Wulf 2018). It is also still poorly understood how RNA polymerase processivity is maintained on the long arrays of tandem repeats (Fingerhut and Yamashita 2022).

Via the analysis of RNA-seq profiles along the complete T2T chicken genome assembly and by RNA-FISH visualisation of nascent transcripts, we demonstrate here that transcription of the whole array of the LL2R tandem repeat is initiated at the 5’UTR of the CR1-H retrotransposable element (Figure 9). Recently, we demonstrated that long terminal repeat (LTR) retrotransposons drive transcription of tandem repeats at subtelomere regions of chicken chromosomes (Krasikova et al. 2024). Together, these results suggest a crucial role of active retrotransposon promoters in the initiation of tandemly repeated non-coding RNA synthesis. It can be suggested that piRNAs targeting active retrotransposable elements may silence transcription of the adjacent tandem repeat arrays (Krasikova et al. 2024).

**Figure 9.**
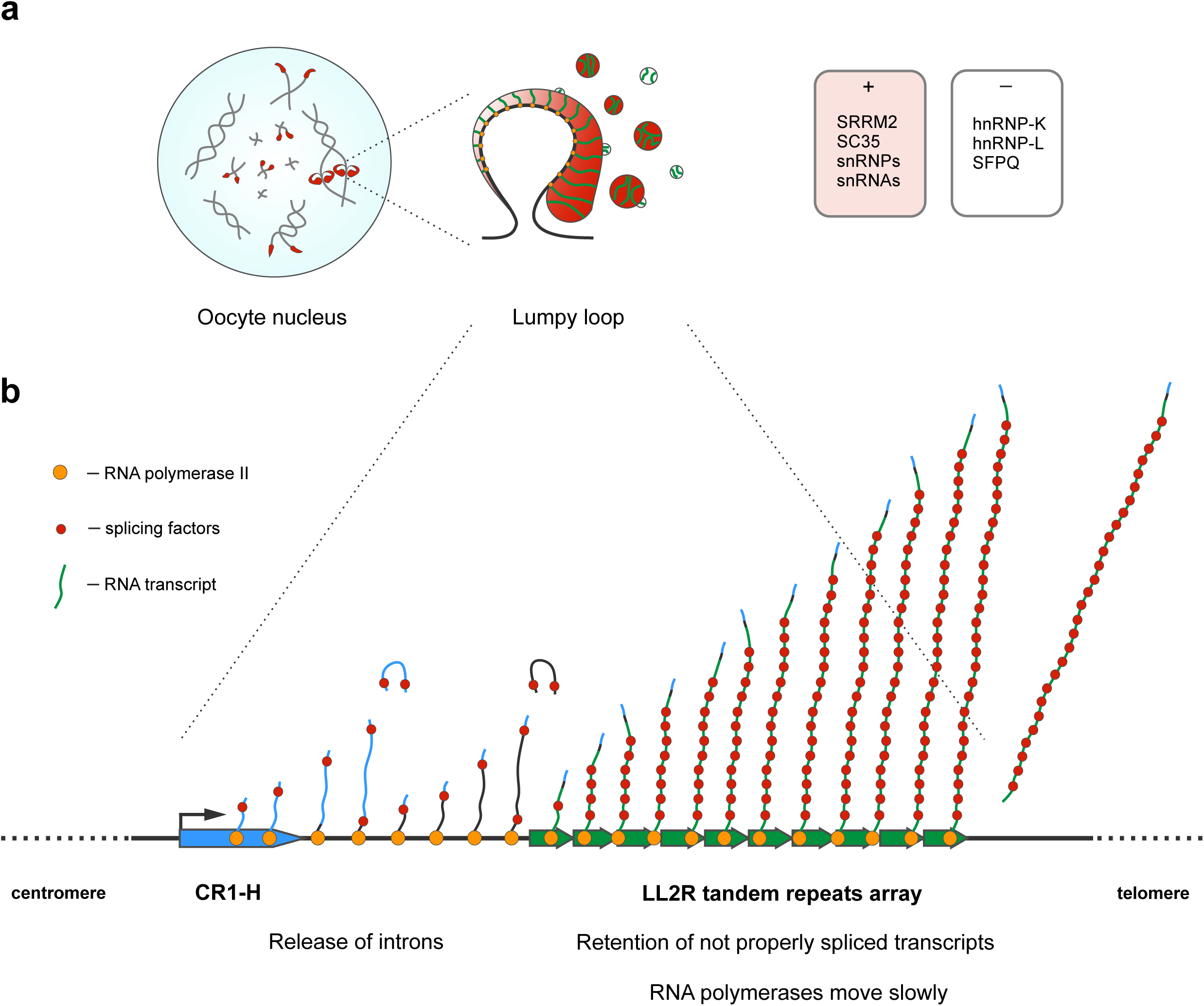
An architectural tandem repeat containing RNA seeds nuclear condensate formation via selective RNA-binding protein (RBP) recruitment and delayed splicing. **a** – Retention of many copies of nascent LL2R repeat containing long non-coding RNAs (green) at the site of transcription driving spatial enrichment of a specific set of splicing factors (red) in the membraneless nuclear domain. Multiple sets of core splicing factors, including serine/arginine-rich proteins SRRM2 and SC35 that contain intrinsically disordered regions, co-transcriptionally associate with LL2R transcripts. At the same time, the RNP-matrix of the LL2R repeat transcription unit (so-called ‘lumpy loop’) lacks hnRNP L, hnRNP K, and SFPQ, widely distributed along normal transcription loops that bear nascent gene transcripts. Partially processed LL2R repeat-containing transcripts form non-diffusible RNP-aggregates enriched by splicing factors that are distributed around LL2R transcription loops and could serve as a site of post-transcriptional splicing. In avian oocyte nuclei at the stage of hypertranscription, splicing factors accumulate within ‘lumpy loops’ and in giant terminal RNP aggregates at the ends of the chromosomes. **b** – LL2R tandem repeat showing RNA transcription dynamics and co-transcriptional processing. Transcription starts at the CR1H retrotransposon (blue) promoter downstream of the LL2R cluster and continues read-through the LL2R tandem repeat array (∼160 kb) that is transcribed very slowly by a convoy of RNA-polymerases with Ser5P CTD (orange circles). Part of the transcript from the CR1-H retrotransposon at the beginning of the transcription unit undergoes normal splicing, following the LL2R repeat containing RNA that binds spliceosome components but is unable to be spliced. Excess of predicted donor splice sites in LL2R repeat-containing RNA leads to deficiency of co-transcriptional splicing of nascent LL2R repeat transcripts, stalling of elongating RNA-polymerases at LL2R transcription units and delay in 3’ end cleavage.

Previously, we found that the most abundant piRNA species in chicken oocyte cytoplasm target CR1 elements (Krasikova et al. 2024) and thus can regulate transcriptional activity of the neighboring LL2R repeat array after fertilization.

Transcription loops of LL2R repeat are covered by a convoy of RNA polymerases with CTD phosphorylated at serine 5 (Ser5) along their entire length. RNA polymerase II CTD Ser5 phosphorylation (Ser5P) peaks in the early elongation phase and at actively spliced exons (Peck et al. 2019). Specifically, catalytically active spliceosomes stably associate with the Ser5P CTD of RNA polymerase II during elongation and co-transcriptional splicing (Nojima et al. 2018). However, the LL2R repeat is transcribed by RNA-polymerase II at a rate lower than in the other lateral loops of lampbrush chromosomes. Extending transcripts of the LL2R repeat do not leave the site of transcription shortly and stay attached to RNA polymerases for a long time. Thus, the LL2R transcription loop demonstrates significant stalling of RNA polymerase II complexes by microscopy-based methods. This stalling of elongating RNA polymerases at LL2R transcription units may be caused by a deficiency of co-transcriptional splicing of nascent LL2R repeat transcripts. On the other hand, 41 bp tandem repeats PO41 and CNM in chicken growing oocytes demonstrate a transcription rate similar to other lampbrush lateral loops (BrUTP being incorporated in 15 minutes after injection) (Deryusheva et al. 2007). Furthermore, nascent PO41 and CNM RNPs with C-rich RNA lack core splicing snRNPs but accumulate hnRNP K. Consequently, we conclude that the LL2R repeat array is transcribed very slowly but forms a loop as it is densely covered by elongating RNA polymerases (Figure 9).

Proper transcription by RNA polymerase II and nascent RNA processing are coupled to ensure successful 5’ end capping, splicing, and 3’ end processing of protein-coding and long non-coding RNA species (Herzel et al. 2017; Peck et al. 2019; Wan et al. 2021). In contrast to protein-coding genes, the LL2R transcript has an excess of the predicted donor compared to acceptor splice sites that may limit the efficiency of its co-transcriptional splicing. Thus, the large size of the RNP-matrix of the LL2R repeat transcription unit is likely due to the accumulation of splicing factors in association with incompletely spliced or completely unspliced nascent LL2R repeat transcripts (Figure 9). Long-read sequencing of nascent RNA demonstrated that the efficiency of co-transcriptional splicing influences the efficiency of 3’ end cleavage (Reimer et al. 2021). In particular, unspliced transcripts demonstrate poor cleavage at gene ends. Moreover, introns with weak 3’ splice sites induce stalling of the splicing reaction (Reimer et al. 2021). Thus, we suggest that 3’ end cleavage of nascent LL2R tandem-repeat containing RNA is unproductive due to incomplete splicing. We also suggest that an excess of the predicted donor splice sites in the LL2R repeat-containing RNA may recruit multiple U1 snRNPs, which are known to repress 3’-end cleavage (Shine et al. 2024).

We further suggest that the transcription unit containing the whole LL2R repeat array spans ∼160 kb due to depletion in polyadenylation signals. At the same time, a fraction of poly(A) LL2R repeat-containing RNA has been revealed in the oocyte nucleus by RNA-seq. Moreover, LL2R transcripts released from template DNA were detected in nucleoplasmic non-diffusable RNP-aggregates and RNP-globules around the LL2R locus (Figure 9). Non-diffusable RNP-aggregates could serve as sites for post-transcriptional splicing of LL2R repeat-containing RNA. Non-diffusable RNP-aggregates could also sequester incompletely spliced LL2R repeat-containing transcripts.

### Mechanism of locus-specific RNP-condensate formation

Biogenesis of many membraneless nuclear compartments is associated with transcriptional activity of certain genomic loci, so that an increase in the local RNA concentration occurs due to intensive transcription. Many copies of an RNA accumulating in proximity to their transcriptional locus can then seed nuclear domain formation (Quinodoz et al. 2021).

Nascent LL2R repeat transcripts are retained at the site of their transcription for a long time, followed by co-transcriptional assembly of nascent RNPs that lack hnRNP L, hnRNP K, and SFPQ (Figure 9). hnRNP L is involved in the export of mRNA into cytoplasm (Dreyfuss et al. 2002). The absence of hnRNP L at certain chromosomal loci may indicate a site of nuclear transcript retention.

LL2R repeat-containing long non-coding RNA functions as a scaffold for multivalent binding of splicing factors guided by its RNA sequence. In particular, LL2R repeat-containing RNA induces the recruitment of a high concentration of RBPs containing IDRs, such as serine/arginine-rich splicing factors SC35 and SRRM2, that can form biomolecular condensates. It was established that IDRs of SRRM2 can mediate the formation of nuclear condensates recruiting splicing factors that modulate SRRM2-mediated alternative splicing (Xu et al. 2022).

Repetitive elements within nuclear retained RNA molecules may be themselves essential in RNA-induced phase separation (Zeng et al. 2022). We found that LL2R RNA comprises short, partly complementary regions, which can potentially form interconnected networks of RNA molecules via partial base-pairing (RNA percolation), facilitating biomolecular condensate formation (Wadsworth et al. 2024). Besides, G-quadruplexes can be predictably formed by LL2R repeat-containing RNA. It is known that G-quadruplexes also undergo percolation within lncRNA-driven condensates (Wadsworth et al. 2024).

Non-diffusible RNP-aggregates that comprise LL2R repeat-containing RNA and associated RBPs are also distributed around LL2R repeat transcription loops in the nucleoplasm. In the lampbrush stage, oocyte nucleus nucleoplasm is enriched in monomeric and oligomeric forms of nuclear actin in a colloidal sol-like state (Maslova and Krasikova 2012). Thus, LL2R RNA-containing RNP-aggregates are suspended in this nucleoplasmic sol.

Transcription loops of the LL2R repeat represent an example of a widespread phenomenon of ‘Sequentially Labeling Loops’

Transcription loops for LL2R repeat array (‘lumpy loops’) represent an example of so-called ‘Sequentially Labeling Loops’ (SLLs) described on lampbrush chromosomes of many amphibian and bird species (Krasikova and Kulikova 2019a). Similar to the ‘lumpy loops’ on chicken lampbrush chromosome 2, individual ‘Sequentially labeling loops’ on lampbrush chromosomes of Urodeles incorporate RNA precursors very slowly, in contrast to thousands of normal transcription loops. ‘Sequentially labeling loops’ include giant granular loops on lampbrush chromosome 12 of Triturus cristatus (Gall and Callan 1962), one of giant fusing loops on chromosome 10 of T. marmoratus (Nardi et al. 1972), giant lumpy loops on lampbrush chromosome 11 of Notophthalmus viridescens (Roth and Gall 1987), globular loops on lampbrush chromosomes of Pleurodeles waltl (Penrad-Mobayed et al. 1986;

Angelier et al. 1990), and special loops on chromosome 3 of Xenopus laevis (Sallacz and Jantsch 2005). Slow rate or even arrest of RNA polymerase movement along the transcription units within SLLs has been proposed (Hartley and Callan 1978; Penrad-Mobayed et al. 1986). Before our study, the loci of SLLs formation and their nascent RNA content remained unknown. Thus, our study represents the first identification of a nascent ’lumpy loop’ or ’sequentially labeling loop’ transcript. The identified LL2R repeat is widespread in the order Galliformes, indicating its functional significance.

‘Lumpy loops’ or other landmark loops with a similar appearance of the RNP-matrix on lampbrush chromosomes of fishes and frogs often accumulate splicing factors (Roth et al. 1990; Eckmann and Jantsch 1999; Sallacz and Jantsch 2005; Dedukh et al. 2013; Dedukh et al. 2025). We propose that nuclear domains enriched in splicing factors in many cases represent locus-specific transcriptional units accumulating distinct sets of RBPs associated with nascent transcripts, rather than free nucleoplasmic nuclear speckles. At the same time, the set of RBP seem to be varied among SLLs. For example, SLLs on lampbrush chromosome 11 of N. viriridescens lack snRNAs involved in splicing and other spliceosome components (Roth and Gall 1987; Wu et al. 1991). In Drosophila spermatocytes, the specific morphology of the nascent RNP-matrix of lampbrush-like loops formed by genes with gigantic introns on the Y chromosome is determined by the repeating RNA sequence within the introns (satellite DNA transcripts) and specific RBPs (Fingerhut et al. 2019; Fingerhut and Yamashita 2022).

### Functions of locus-specific RNP condensates

Biomolecular condensates can sequester or inactivate splicing factors, serve as dynamic storage for multiple mRNAs and RBPs, and regulate alternative splicing (Giudice and Jiang 2024). Appearance of tandem repeats within arcRNAs multiplies the number of binding sites for the same set of RBPs. LL2R repeat-containing RNA can be involved in the regulation of splicing via deposition of certain sets of core splicing factors in an immobile but dynamic nuclear domain associated with a specific chromosomal locus (Figure 9). ‘Lumpy loops’ at the loci of nascent LL2R repeat RNA synthesis concentrate factors required for splicing, but apparently lack any client RNA molecules due to the absence of polyadenylated RNA. The chicken oocyte nucleus at the stage of hypertranscription lacks any nucleoplasmic nuclear speckles (Krasikova et al. 2012), and efficient splicing occurs presumably co-transcriptionally at nascent gene transcripts (Krasikova et al. 2024).

LL2R repeat transcription loops that concentrate splicing factors are closely related to nuclear stress bodies in human somatic cells. Nuclear stress bodies (nSBs) form around human pericentromere satellite III (HSATIII) transcripts upon stress or pathological conditions and serve as a platform to sequester hundreds of RBPs (Ninomiya et al. 2023). HSATIII lncRNAs are transcribed by RNA polymerase II (Jolly et al. 2004) and recruit a set of RBPs, including SR-splicing factors (SRSF1/SF2-ASF, SRSF7, SRSF9) (Denegri et al. 2001) and hnRNPs M, A1, and H1 (Chiodi et al. 2000; Aly et al. 2019). During stress recovery, nSBs recruit CDC-like kinase 1 (CLK1), thus facilitating SR-protein phosphorylation and subsequently promoting intron retention in a wide range of pre-mRNAs (Ninomiya et al. 2020). In contrast to nSBs, transcription loops for the LL2R repeat array appear during normal oocyte development.

Oocytes may also use phase transition of RBPs to control their solubility over an extended period of time, but begin their dissociation during maturation (so-called ‘arrested’ condensates) (Pek 2025). We propose that the transcription loops of the LL2R tandem repeat array on avian lampbrush chromosomes also act as a depot for splicing factors in the oocyte nucleus. Such a depot probably regulate concentration of a set of RBPs during oocyte growth by their sequestering and subsequent release.

## Conclusions

The combination of bioinformatic genome and oocyte nucleus transcriptome analysis and ultrastructural cytogenetic imaging of highly extended lampbrush chromosomes allowed us to identify individual transcription units of a novel LL2R tandem repeat at specific chromosomal loci and provide a detailed view of the LL2R repeat transcription and transcript processing, not achievable in the interphase nucleus. We demonstrated that an active retrotransposable element drives transcription of the LL2R tandem repeat and thus participates in loci-specific nuclear domain formation. Tandem repeat transcripts nucleate membraneless nuclear compartment formation serving as an architectural RNA by containing excess/deficiency of particular RBP sites, thus leading to concentration/depletion of the particular RBPs at a particular chromosomal locus. The LL2R repeat-containing arcRNA contains an excess of donor splice sites and thus recruits specific sets of multiple splicing factors that contain IDRs and can form nuclear condensates. Due to the deficiency of acceptor splice sites, impaired splicing results in a very low rate of transcription and retention of strand-specific nascent LL2R repeat-containing RNA with associated RBPs on the transcription loops.

## Acknowledgements

The research was performed using the equipment of the Resource Centers ‘Molecular and Cell Technologies’, ‘Interdisciplinary Resource Centre for Nanotechnology’, ‘Chromas Core Facility’, and ‘Centre for Microscopy and Microanalysis’ (Saint-Petersburg State University).

## Author contributions

AK conceived and designed the study. AK and AF conducted the bioinformatic analysis and designed FISH probes, AZ performed molecular cloning, AK, AZ and TK isolated lampbrush chromosomes, AK performed LL2R FISH mapping experiments, AZ performed ribonuclease treatment assay, AK stained the isolated oocyte nuclei followed by confocal laser scanning microscopy, AK, AF and TK performed chromosome immunostaining experiments, AF made immuno-FISH experiments, TK made BrUTP microinjections, AK and TK performed AFM and SEM, VS performed 3D-SIM. AK analyzed the data and wrote the manuscript. All authors provided critical feedback and approved the final version of the manuscript.

## Supplementary Materials

Supplementary Materials contain 8 additional figures.

