## Supplementary Materials for "Nascent transcripts of LL2R tandem repeat nucleate locus-specific RNP-condensates recruiting splicing factors"

**Supplementary Figure S1. 'Lumpy loops' on chicken lampbrush chromosome 2 visualized by light and atomic force microscopy (AFM). a** – Lampbrush chromosome 2 stained with DAPI (greyscale); **a'** – Corresponding phase contrast image. Arrows indicate 'lumpy loops'. Scale bar = 10 µm. **b** – AFM of 'lumpy loops' framed on **a'**; **c** – 3D reconstruction of 'lumpy loops' shown in **b**.

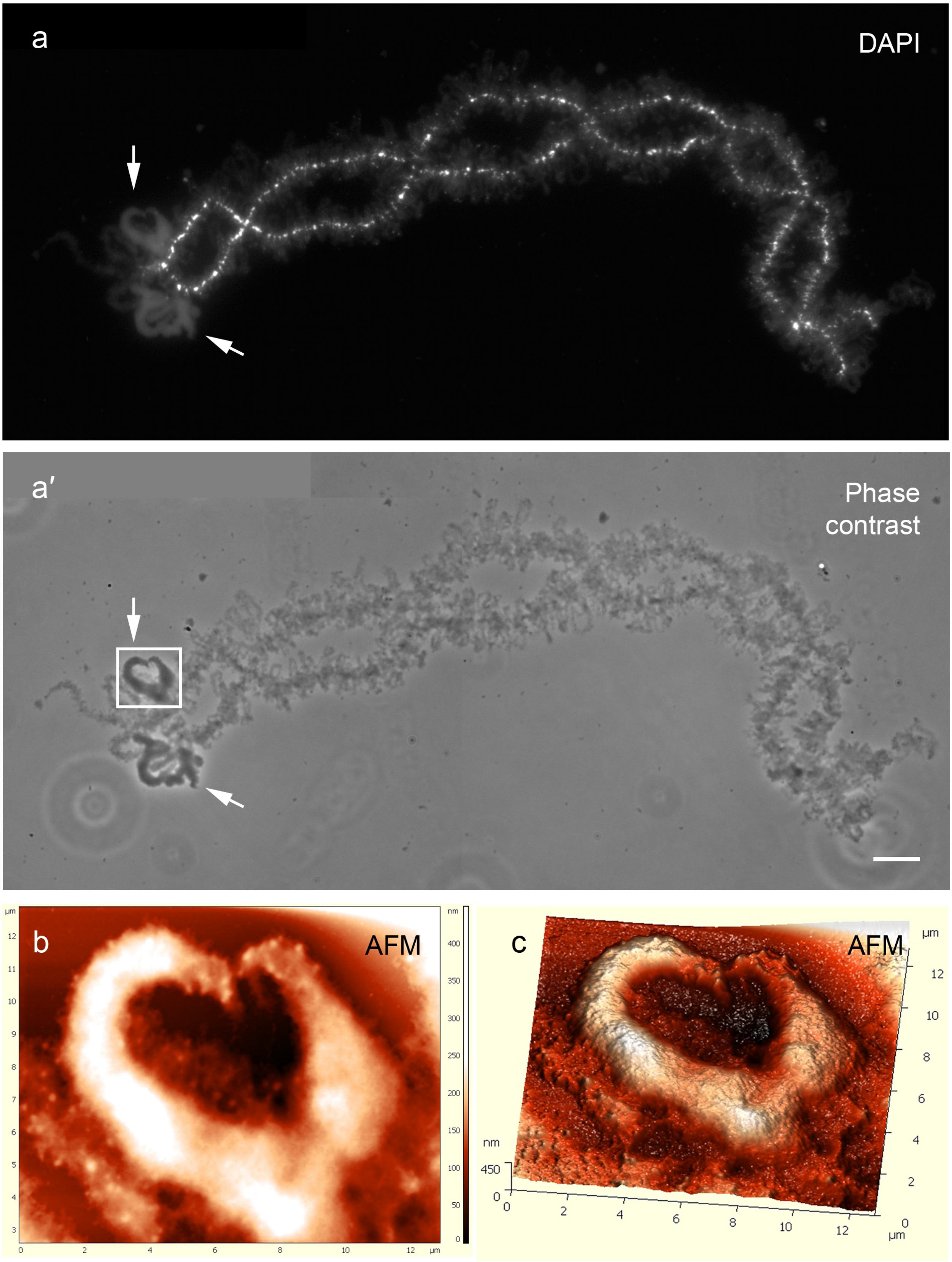

**Supplementary Figure S2. Identification of LL2R-like repeats in the turkey genome. a** – Overview of the turkey (*M. gallopavo*) chromosome 3 region containing LL2R-like repeats (according to MGAL_WU_HG_1.0 genome assembly). **b** – Enlarged region showing the density of predicted splice sites and polyadenylation signals across the LL2R-like repeat array. Nearly full-length CR1-H element (corresponding to positions 4-323, 1800-2867, 3408-4799 of the consensus) precedes the LL2R-like repeat array. The Masked repeats track visualizes the positions of repeats from the Repbase library of chicken repeats. Individual predicted donor and acceptor splice sites and polyadenylation signals are shown by vertical bars. The graphs below show the average (in a 2 kb window) density of predicted splice sites. Vertical dotted lines indicate the position of the boundaries of the LL2R-like repeat array. **c** –Turkey chromosome 3 stained with Y12 antibodies against the Sm epitope of snRNPs (green); **c'** – corresponding phase contrast image. Chromosomes are counterstained with DAPI (blue). Scale bar = 10 µm.

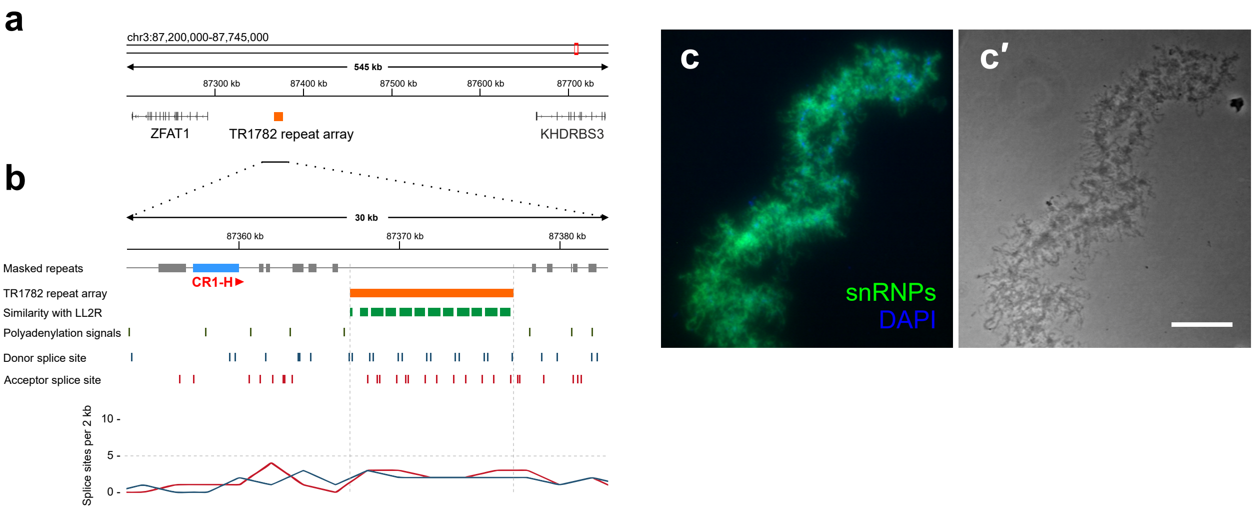

**Supplementary Figure S3. LL2R transcripts sustained after treatment with RNases H, III and R.** RNA-FISH with the LL2R probe (green) and BAC-clone WAG-13J24 (red) after treatment with RNase H (**a**), RNase III (**b**), RNase R (**c**), and without any RNase treatment (**d**). Chromosomes are stained with DAPI (blue). **a'-d'** – Corresponding phase contrast images. Arrows indicate 'lumpy loops' containing transcripts of LL2R repeats on chicken lampbrush chromosome 2. Scale bars = 10 μm.

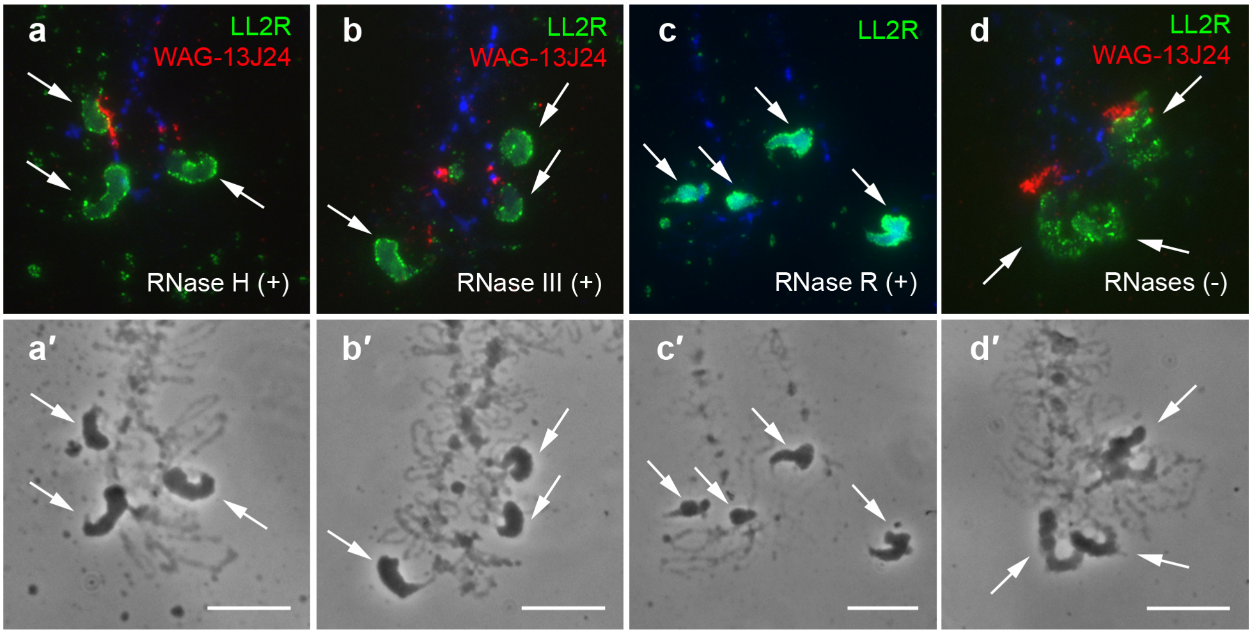

**Supplementary Figure S4. The RNP-matrix of 'lumpy loops' on lampbrush chromosome 2 lacks polyadenylated RNA.** RNA FISH with oligo-dT(30) (green), chromosomes are stained with DAPI (blue). **a** – Fragment of lampbrush chromosome 2 with 'lumpy loops' (arrows) and oligo-dT(30) positive bow-like loops (pink arrowheads); **b** – A microchromosome bearing poly(A)-rich RNP aggregates at its terminal region (GITERA, white arrowheads). **a', b'** – Corresponding images with signal from oligo-dT(30) only. **a'', b''** – Corresponding phase contrast images. Scale bars = 10 µm.

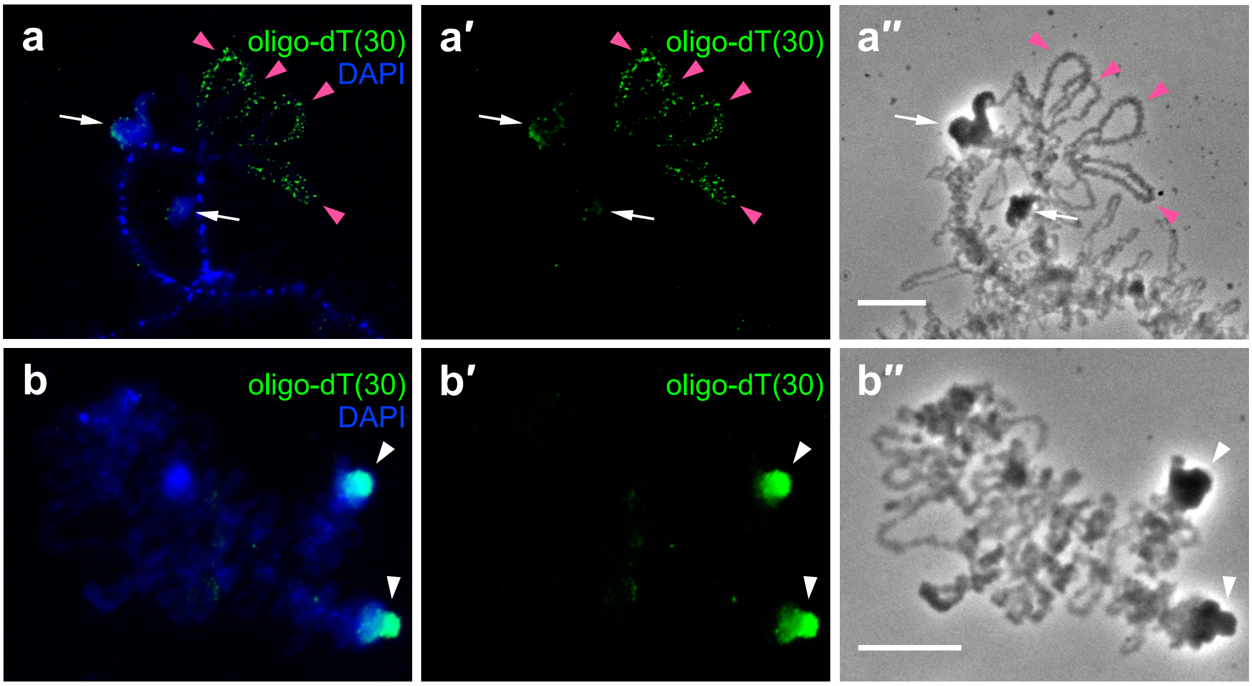

**Supplementary Figure S5. Potential RNA-RNA interactions and predicted formation of G-quadruplexes by LL2R repeat-containing RNA. a** – Potential RNA-RNA interactions within LL2R repeat-containing RNA (TR904 consensus) predicted with IntaRNA program; **b** – Potential formation of G-quadruplexes by LL2R repeat-containing RNA (TR904 consensus) predicted by QGRS Mapper. **c** – Position of oligonucleotide probes according to donor splice sites and potential RNA-RNA interaction regions (TR904 consensus).

### a RNA-RNA interactions

```
          769                                                                                                                    870
             |                                                                                                                   |
5'-UGG..CAUUU     C           GG                  AAAAAUG        CAUGG   U       GG    C    A            AAUG   A        GG      GAAGC..GGG-3'
             CUGAC   AGAGUUUCC      GACGGA  UGUGAA       GCCUCAGG     AUA GAUGGAA  AACG GGUG GCA AGGU  CC     AG GAGGGCAG    AGGU
             |||||   |||||||||      ||||||  ||||||       ||||||||     ||| |||||||  |||| |||| ||| ||||  ||     || ||||||||    ||||
             GACUG   UCUCGAGGG      CUGUCU  ACAUUU       CGGGGUCC     UGU CUACUUU  UUGC CCAC CGU UUUA  GG     UC CUCUCGUC    UCCA
3'-GGG..UGUGU     UUU         AAGUUU      UU      AAAGU          UUUU    C                  C   A    CU  AAUGG           AGCG    GAGGG..GGU-5'
             |                                                                                                                   |
          230                                                                                                                    122
```

| Target | Start(T) | End(T) | Query | Start(Q) | End(Q) | Energy |
|---|---|---|---|---|---|---|
| LL2R | 769 | 870 | LL2R | 122 | 230 | -30.68 |

### b QGRS sequences found

| Position | Length | QGRS | G-Score |
|---|---|---|---|
| 2 | 23 | **GG**T**GG**CTGCTCGTCCGT**GG**TT**GG** | 10 |
| 42 | 28 | **GG**GT**GG**CAGAT**GG**CCATGCATGTGTT**GG** | 10 |
| 409 | 21 | **GG**GCA**GG**GT**GG**TGAAGCCC**GG** | 15 |
| 441 | 13 | **GG**AT**GG**AT**GG**T**GG** | 20 |
| 543 | 30 | **GG**AAAGAGTAA**GG**TAATG**GG**AGACCTGC**GG** | 17 |
| 724 | 28 | **GG**TATAGTGATGAATTTGCT**GG**T**GG**T**GG** | 4 |
| 804 | 23 | **GG**CCTCA**GG**CAT**GG**ATATGAT**GG** | 17 |
| 860 | 20 | **GG**CA**GG**GA**GG**TGAAGCCA**GG** | 15 |

### c Position of oligonucleotide probes

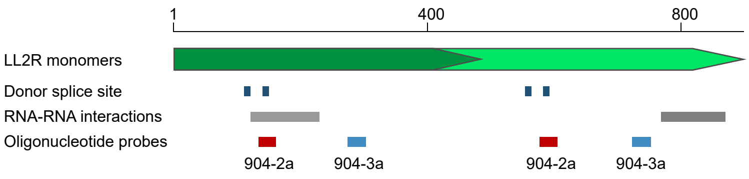

**Supplementary Figure S6. 3D-SIM showing the distribution of RNA polymerase II and double-stranded DNA on the 'lumpy loops' bearing nascent transcripts of LL2R repeats. a**, **b** – Immunodetection of RNA polymerase II phosphorylated at serine 5 of the CTD repeat (green) followed by RNA-FISH with the PCR probe to LL2R repeats (red); **a'**, **a''** – Enlarged fragment of **a**. **c, d** – Immunodetection of double-stranded DNA (green) followed by RNA-FISH with oligonucleotide probe 904-3a to LL2R repeats (red); **c'**, **c''** – enlarged fragment of **c**. Arrows indicate 'lumpy loops', arrowheads indicate RNP aggregates, and orange curved arrows indicate the direction of transcription. Chromosomes are counterstained with DAPI (blue). Scale bars = 5 µm.

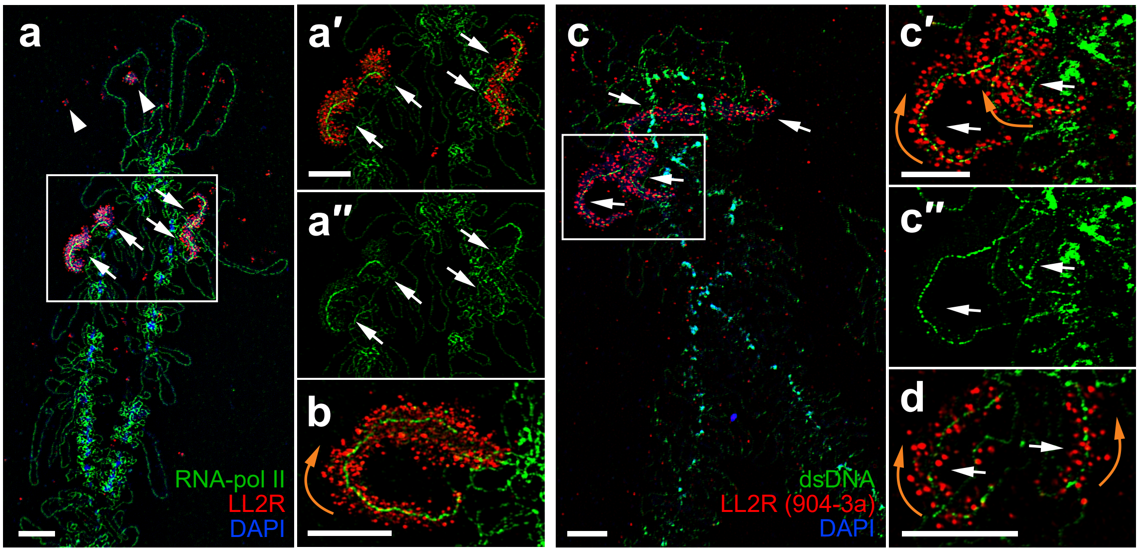

**Supplementary Figure S7. 3D-SIM showing the colocalization of nascent LL2R repeat-containing RNA and splicing factors on the 'lumpy loops' of chicken lampbrush chromosome 2.** Immunodetection of TMG-capped snRNAs with K121 antibodies (**a**, **a'**, **b**, **b'**), Sm proteins of snRNPs with Y12 antibodies (**c**, **c'**) or factors SRRM2 and SC35 (**d**, **d'**) (green) followed by RNA-FISH with the PCR probe to LL2R repeat with (**a**, **a'**) or without (**b**, **b'**, **c**, **c'**, **d**, **d'**) Triton X-100 pretreatment. Pre-treatment with Triton X-100 is necessary for the permeability of the PCR-based probe into the RNP-matrix of 'lumpy loops'. Arrows indicate 'lumpy loops', arrowheads indicate RNP-aggregates. Chromosomes are counterstained with DAPI (blue). Scale bars = 5 µm.

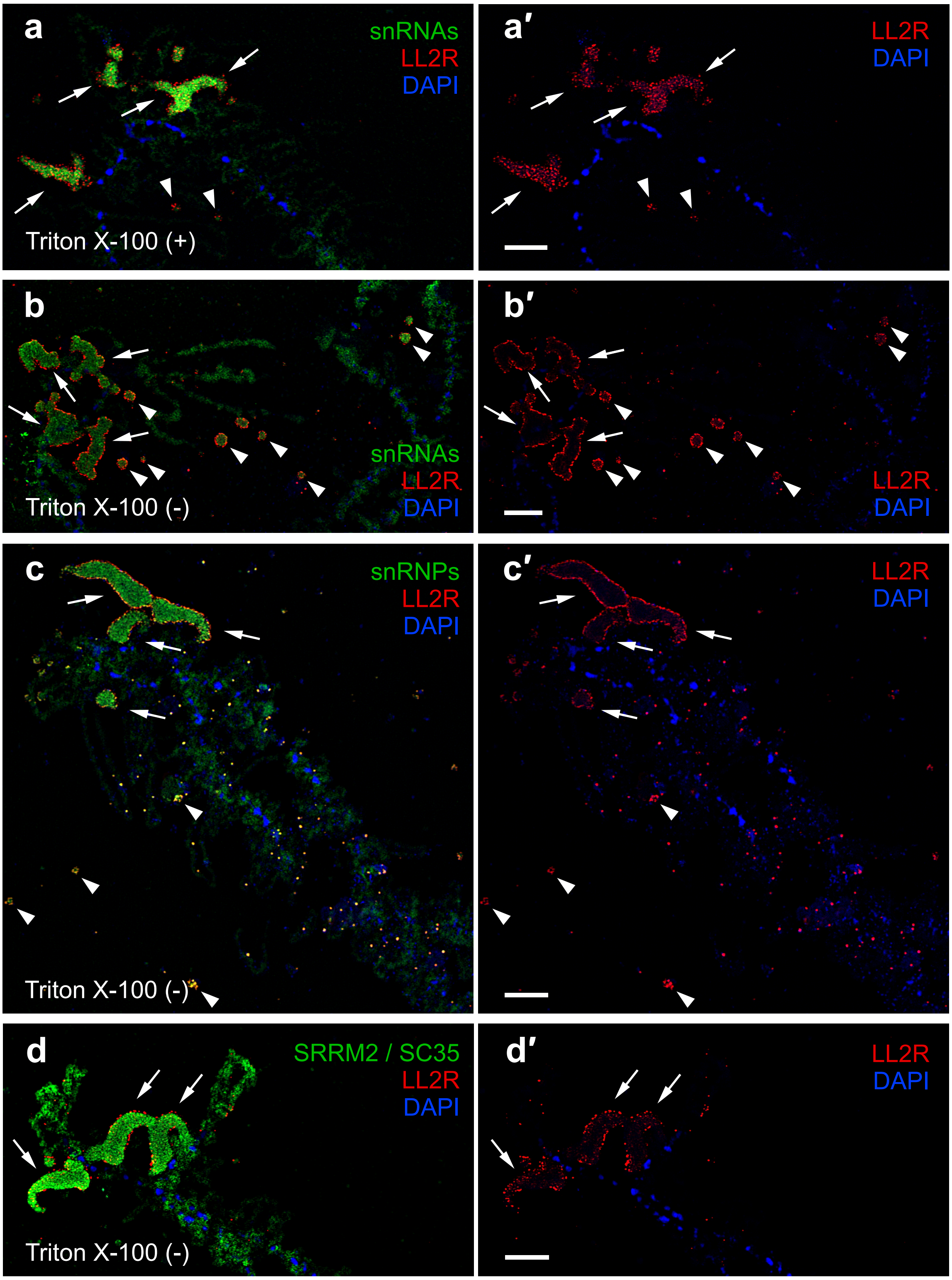

**Supplementary Figure S8. 'Lumpy loops' on lampbrush chromosome 2 accumulate phosphorylated SC35.** Immunodetection of hypophosphorylated SC35 (green) after treatment with calf intestinal phosphatase (CIAP) (**a**) and without treatment (**b**). Chromosomes are counterstained with DAPI (blue). **a'**, **b'** – corresponding phase contrast images. Arrows indicated 'lumpy loops' on lampbrush chromosome 2. Scale bars = 10 µm.

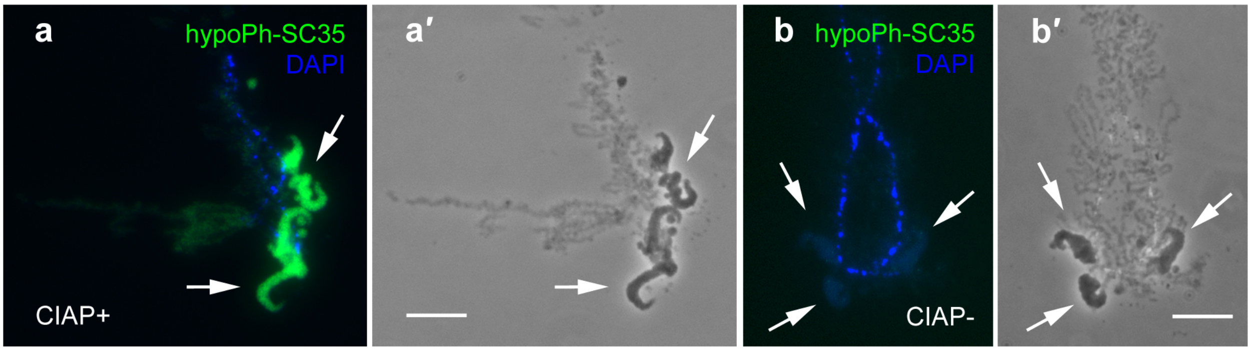
